# Local sleep in wake is widespread across the rat cortex and tracks sleep need

**DOI:** 10.64898/2026.11.22.768422

**Authors:** Graham Findlay, Matias Lorenzo Cavelli, Tom Bugnon, William Marshall, Giulio Tononi, Chiara Cirelli

## Abstract

During non-rapid eye movement (NREM) sleep, cortical neurons alternate between ON periods of firing and OFF periods of silence, the latter accounting for the slow waves of the EEG. Converging evidence indicates that OFF periods can also occur locally during extended wake (“local sleep”), at a behavioral cost, but whether this phenomenon is ubiquitous across the cortex, whether some areas are more resilient than others, and whether it can also provide any benefit, remain open questions. Here we recorded from 11 cortical areas (and claustrum) in 15 rats using high-density Neuropixels probes and detected OFF periods with a novel method that captures spatially restricted and irregularly shaped events mischaracterized or missed by traditional methods. We found OFF periods during extended wake in every region tested, with no evidence that any area is spared. Over the course of sleep deprivation, OFF periods became more frequent, longer, larger, and quieter, and their features correlated with delta power on a per-event basis in both wake and NREM sleep, consistent with the notion that they reflect accumulating sleep need. The extent to which OFF periods overlapped across areas also increased across sleep deprivation. Finally, the rate of medium and large OFF periods during sleep deprivation was negatively correlated with their rebound in subsequent recovery sleep, consistent with the possibility that local sleep in wake partially discharges sleep need before the animal falls asleep.

**Statement of Significance:** Prolonged wakefulness drives small groups of cortical neurons into brief sleep-like states even as the animal remains awake and behaving. This “local sleep” in wake is thought to underlie the errors and slowed responses of sleep deprivation, but how widespread it is, and whether it helps, remains unclear. Using high-density recordings across many cortical regions in the rat and a novel detection method sensitive to small, irregular events, we show that local sleep is not confined to a few vulnerable areas but occurs throughout the cortex, growing with accumulating sleep need. We also find that these neuronal lapses may begin to relieve that need before sleep takes over, suggesting that they may represent an early, piecemeal form of recovery.

## Introduction

In the corticothalamic system, pyramidal neurons tend to be persistently active during wake, even more so during learning [1, 2]. During non-rapid eye movement (NREM) sleep instead, neurons alternate every half a second or so between ON periods of tonic activity, when they fire at rates similar to those in wake, and OFF periods of silence, when they are hyperpolarized and stop firing altogether [3]. ON and OFF periods occur more or less synchronously in many neurons [3], accounting for the slow waves that can be recorded using electroencephalography (EEG). Slow wave (delta) activity, a combined measure of the number and amplitude of slow waves, is defined as the EEG power in the low frequency range of 0.5 to 4 Hz.

When waking is prolonged beyond its physiological duration (∼16 hours in humans), “microsleep” episodes lasting a few seconds can happen, during which we become disconnected from the environment and muscle tone drops. Like longer periods of NREM sleep, microsleeps reflect the occurrence of ON/OFF patterns of activity across the whole thalamocortical circuit, can be detected by the scalp EEG, and measured as delta activity. However, there is converging evidence that even when responsive to stimuli and engaged in behavior, subjects can undergo episodes of “local sleep” despite an EEG that is typical of waking. This phenomenon is documented in both rodents and humans. Specifically, unit recordings in rats showed that, after a few hours of sleep deprivation, small subsets of cortical neurons undergo OFF periods as they usually do in NREM sleep, but they do so intermittently, at changing locations, while the animal is awake and behaving [4]. Because they are local, these OFF periods do not result in slow waves detectable by the standard scalp EEG, but only in the local field potential, where they appear as single, isolated waves in the 2 to 6 Hz range [4]. Local sleep can be inferred in awake humans by employing high-density EEG (hdEEG, 256 channels) that can resolve small, local theta (5–8 Hz) waves on the scalp [5–7]. In sleep deprived humans the local increase in theta waves detected with hdEEG correlates with impaired performance [5]. Intracranial recordings in the temporal lobe of epileptic patients also show that sleep deprivation leads to “neuronal lapses” associated with cognitive lapses [6]. Local slow waves can also occur in non-sleep deprived humans after intense learning, when they are associated with impaired performance [7, 8]. The latter can be reversed by a nap but not by a wake period of similar duration [7].

Despite the potential broad consequences for performance, many aspects of local sleep in wake remain poorly understood. Here we address three questions in freely moving rats chronically implanted with Neuropixels probes, which allow dense sampling of cortical activity [9]. First, is local sleep ubiquitous across the cortex, or are some areas spared? Second, do the OFF periods of extended wake, including small and irregularly shaped events that conventional detection methods are not designed to capture, track sleep need in their number and size and in their relation to delta activity? Third, despite its behavioral cost, does local sleep help discharge the sleep pressure accumulated during extended waking? To address these questions we first describe a detection method that, rather than classifying a recorded population as ON or OFF together on the basis of its pooled activity, resolves OFF periods in depth as well as in time, so that spatially restricted and irregularly shaped events are counted as well.

## Methods

Most (13/15) of the animals described in the current study were used in a recently published paper analyzing the relationship between cortical and hippocampal sleep [10], and the same protocols were followed for the remaining (2/15) animals. Surgery, experimental procedures and design, histology, electrophysiological recordings, local field potential (LFP) preprocessing, spike sorting, and sleep scoring are therefore described in [10] and are summarized below only where necessary. Here we focus on the aspects of the analysis that are unique to the present work.

### Overview of the analysis

We detected OFF periods on each probe from the envelope of multiunit activity (“Morphological OFF period detection”). Each OFF period was described by its duration, span, area and residual multiunit activity (how “silent” the OFF was), and these descriptors were summarized per subject, cortical structure and experimental condition. We then asked four questions, each with its own subsection below: whether these summaries change with time awake and asleep across six conditions, after brief sleep intrusions had been excluded from wake (“Sleep intrusion scoring”, “Definition of statistical conditions”, “Statistical analysis”); whether and how OFF-period features track delta power, event by event and epoch by epoch (“Per-event correlations between OFF properties and instantaneous delta power”, “Epoched partial correlations between OFF area and delta power”); whether OFF periods in different areas coincide more often than chance predicts (“OFF locality/overlap”); and whether the OFF periods expressed during sleep deprivation predict the NREM rebound that follows (“Correlation between sleep deprivation and subsequent NREM rebound”).

### Experimental animals

Adult rats (Sprague Dawley, males, 300–340 g, 2–3 months old; RRID: RGD_734476) were maintained on a 12 h light/12 h dark cycle with food and water available *ad libitum* (21–26 °C, 30–40% relative humidity). Animals were group housed until the time of surgery. All animals were healthy, drug-naive, and were not used in previous procedures. All animal procedures and experimental protocols followed the National Institutes of Health Guide for the Care and Use of Laboratory Animals and were approved by the Institutional Animal Care and Use Committee (IACUC) of the University of Wisconsin-Madison. Animal facilities were inspected and accredited by the Association for Assessment and Accreditation of Laboratory Animal Care (AAALAC).

### Morphological OFF period detection

To resolve OFF periods in depth as well as in time, we detected them as connected regions of low activity in a channel-by-time image of the envelope of high-frequency activity, rather than as intervals of silence in pooled spike times [11]. Both the envelope of high-frequency activity and unsorted spike times are often referred to as “multiunit activity” (MUA) – an unfortunate terminological collision – but we always mean the former. MUA traces were obtained by bandpass filtering the raw SpikeGLX AP-band traces between 300 and 5,000 Hz; phase-shifting to correct for the Neuropixels’ analog-to-digital multiplexing (sampling) delays; automatic detection and interpolation of bad channels; a spatial highpass (‘destriping’) filter to suppress spatially coherent artifacts; full-wave rectification; and antialiased resampling to 500 Hz. These steps were all performed using their respective SpikeInterface v0.104 preprocessing functions with default arguments. At detection time, each channel’s envelope was further smoothed with a Gaussian lowpass filter (20 Hz cutoff, corresponding to a standard deviation of ∼8 ms) followed by a 15-sample (30 ms) temporal median filter (Supplementary Fig. 1a).

Within each structure, channels near its edges were excluded if, on visual inspection of MUA traces across the sleep/wake cycle, a channel appeared “silent” (lacking MUA), or if there was visible bistability but its magnitude was greatly reduced compared to the rest of the structure, suggesting a lack of signal-to-noise ratio rather than a lack of bistability (Supplementary Fig. 2). The remaining contiguous block of channels was used for detection in all states and conditions (median span excluded 310 µm, IQR 0-600 µm). Although exclusions were based on visual inspection, the visually excluded and retained channels separate almost perfectly on MUA envelope standard deviation (median AUC 0.95 during baseline NREM). Because we cannot with certainty say that a lack of bistability on a channel at the margin of a structure is due to a lack of MUA rather than a real lack of bistability, we make no claims about the ubiquity of bistability at the channel level.

The threshold for each channel was a quantile of that channel’s own envelope distribution rather than an absolute value (Supplementary Fig. 1b). It was recomputed for every channel and every ∼5.5-min processing block, so that the same fraction of samples fell below threshold on every channel and in every block. One quantile was specified per structure for NREM sleep and one for wake: the NREM threshold for a block was that quantile of the block’s NREM samples, and the wake threshold was that quantile of its wake samples. Artifactual samples were never eligible to be part of an OFF period and were never included in quantile estimation. We used quantiles because the MUA envelope is not amplitude-normalized and its noise level can drift over the many hours separating the periods we wished to compare, and we recomputed them block by block because a single whole-recording quantile was not stable across blocks (Supplementary Methods). Because the same fraction of samples was marked subthreshold before morphological cleaning (see below), comparisons of OFF duration and area between conditions were not biased by drifts in signal scale; for the same reason, differences in OFF rate and area between conditions reflect how subthreshold samples cluster in space and time, not how many there are (Supplementary Methods). For each structure, the two quantiles were set manually, by subjective visual inspection of the resulting detections across the whole 48 hours, to balance false positives against false negatives; in setting them, we deliberately tolerated false negatives (missed OFF periods) more than false positives (spurious detections). Across structures the median NREM quantile was 12% (median absolute deviation 3%), and the median wake quantile was 0.6% (median absolute deviation 0.3%).

OFF periods were then detected as follows. Each channel’s smoothed envelope was thresholded to produce a binary channel-by-time image of subthreshold samples (Supplementary Fig. 1c), which was cleaned with a sequence of binary morphological operations (Supplementary Fig. 1d): a spatial closing across 4 channels (80 µm) to bridge occasional lower signal-to-noise ratio channels; a temporal opening of 15 samples (30 ms) to remove brief, isolated threshold crossings; a spatial opening across 4 channels (80 µm) to remove events involving too few channels; and a final spatial closing across 3 channels (60 µm) to connect vertically adjacent regions. Candidate OFF periods were defined as the connected components (‘blobs’) of the cleaned binary image (four-connectivity; ‘scipy.ndimage.label’; Supplementary Fig. 1e).

Candidate OFF periods were then admitted by one of two nested size criteria. For each event we measured its vertical span (the depth difference between its deepest and most superficial involved channel), its median duration (the median, across involved channels, of each channel’s subthreshold duration) and its total duration (from the earliest onset to the latest offset across involved channels). A relaxed criterion required a span of at least 100 µm, a median duration of at least 25 ms and a total duration of at least 30 ms; a stringent criterion required a span of at least 200 µm, a median duration of at least 50 ms and a total duration of at least 60 ms (Supplementary Fig. 1f). The stringent duration criteria follow prior work [4, 11]. Because the spatial opening removes any event spanning fewer than 4 channels (a span below 60 µm), 100 µm is the smallest span comfortably above this floor; the stringent span of 200 µm is twice that. The relaxed criteria are half the stringent ones, so as to include events that would conventionally be considered too small. We refer to events admitted by the relaxed but not the stringent criterion as ‘small’, and to events admitted by the stringent criterion as ‘medium/large’ (Supplementary Fig. 1g). No upper bound on duration was imposed.

For each admitted OFF period we extracted descriptors of its temporal and spatial extent (Fig. 1f). In addition to the span and median duration defined above, we computed its area, the number of channel-by-sample pixels in the event’s mask (one pixel is 20 µm by 2 ms), and its median MUA, the median of the rectified, smoothed MUA envelope across all pixels of the event. Median MUA measures the residual multiunit firing during the OFF period (lower values indicating a ‘deeper’, more completely silenced event). To summarize all OFF periods within a condition in a way that is comparable across structures of different span and conditions of different length (late baseline sleep was not exactly 1 h long; see “Definition of statistical conditions” below), we computed the total normalized area (or “total OFFness”): the summed area of all of a condition’s OFF periods, divided by the structure’s maximum possible span and by the duration of the condition. Finally, to measure how synchronously a neural population entered or exited an OFF period, we computed the median absolute deviation of the per-channel onset (or offset) times across the channels the event spans (“onset/offset asynchrony”); larger values denote a more asynchronous edge. Values of onset/offset asynchrony are reported after adjustment for mechanical dependence on span (see “Correcting for mechanical correlation of edge asynchrony and span” in Supplementary Methods).

**Figure 1.**
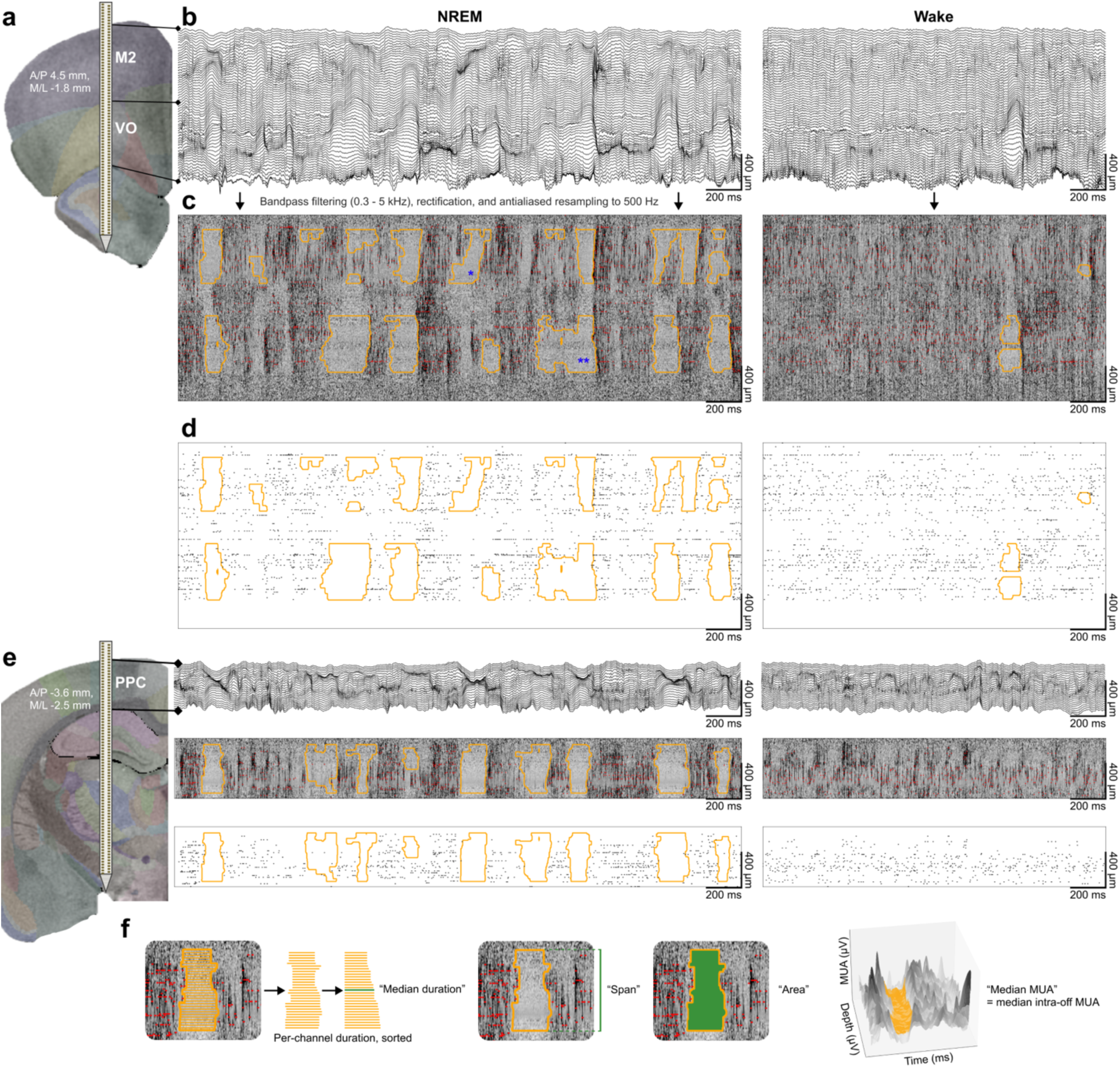
Morphological detection of OFF periods during NREM sleep and wake. **a,** coronal section from the Waxholm Space Rat Brain Atlas, showing implant location of a Neuropixels 1.0 probe in secondary motor cortex (M2) and ventral orbital cortex (VO) of an example subject. A/P coordinates are relative to bregma. **b,** example local field potentials (LFPs) spanning M2 and VO during NREM sleep and wake. **c,** example “multiunit activity” (MUA) traces, obtained by bandpass filtering (0.3-5 kHz), rectifying, and downsampling the wideband signals. Darker shades indicate a higher envelope of high-frequency activity. Morphologically detected OFF periods are outlined in orange (see Methods and Supplementary Fig. 1). Spike sorted unit clusters are overlaid in red but were not used for morphological OFF detection. Even in this ∼2 s example, a diversity of OFF sizes and shapes is apparent. The single blue asterisk highlights an OFF period with “diagonal” onset and offset that start in deep layers and end in superficial layers of cortex. The whole column is silent together for only ∼40 ms, which can cause traditional OFF detection methods to either underestimate the size or miss OFF periods entirely. The double blue asterisk illustrates another OFF period difficult to capture with traditional OFF detection, because whole-column silence comprises only about 1/4 of its total duration. **d,** the same morphologically detected OFF periods (orange) and spike rasters (not used for morphological detection) shown in c, but without underlying MUA traces. **e,** same as (a-d), for a second Neuropixels probe implanted simultaneously in posterior parietal cortex (PPC) of the same rat. **f,** temporal and spatial properties of OFF periods.

### Delta power computation

To relate OFF periods to delta activity, both across conditions (Fig. 2f) and at the time of individual events (Fig. 3), we computed delta power from the LFP of each structure using a filter-Hilbert method, as in our previous work [10]. For each probe, the LFP (sampled at ∼625 Hz) was first bipolar-referenced by subtracting from each contact the signal recorded 200 µm (10 contacts) deeper, which suppresses volume-conducted, spatially broad signals and emphasizes local activity. The bipolar LFP was decimated to 31.25 Hz with zero-phase antialiasing, bandpass filtered in the delta band (0.5–4 Hz) with a zero-phase second-order Butterworth filter, and converted to an analytic signal by the Hilbert transform; instantaneous delta power was taken as the squared amplitude of the analytic signal. A per-structure delta power time series was obtained as the median instantaneous power across all bipolar channels lying entirely within the structure.

**Figure 2.**
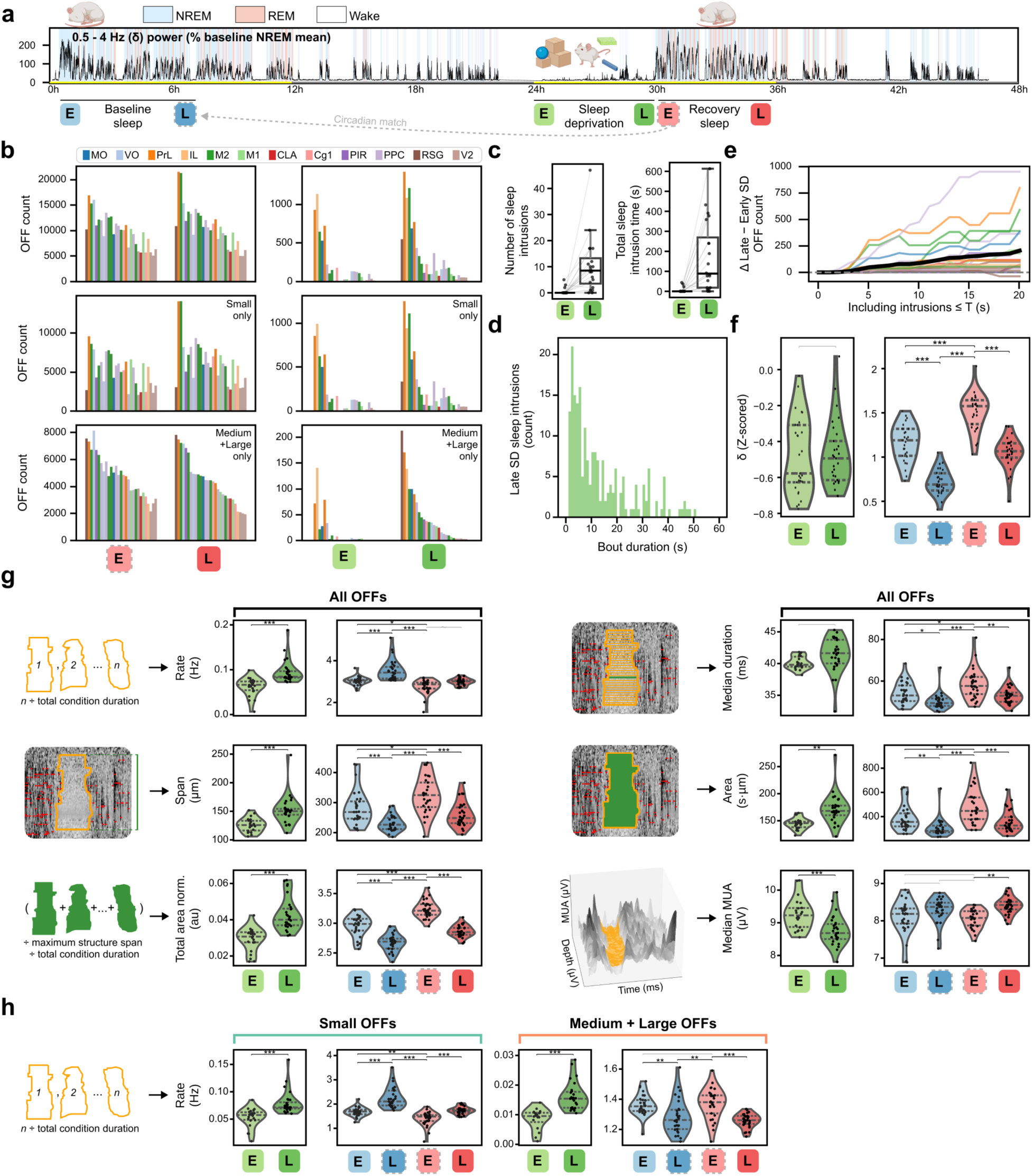
Sleep/wake homeostatic changes in OFF periods. **a,** experimental protocol: 48 h of continuous recording including 6 h of sleep deprivation. Sleep scoring and cortical delta (0.5-4 Hz) are shown for one example rat. Yellow and gray bands along the time axis indicate the light and dark periods, respectively. Colored boxes indicate the six conditions (1 h each, except late baseline sleep; “E” for early, “L” for late) used to assess homeostatic effects in subsequent analyses. Early recovery sleep (light red) is the first cumulative hour of sleep following late sleep deprivation (dark green), and late recovery sleep (dark red) is the final hour of recovery sleep before lights-off. Early baseline sleep (light blue) is the first hour of sleep following baseline lights-on, and late baseline sleep (dark blue) is the sleep that occurred 24 h before early recovery sleep (i.e., at the same circadian time as early recovery sleep; see Methods). **b,** left: change in OFF counts from early to late recovery sleep using 2 sets of criteria to define OFF periods (n=29 unique subject-structure pairs from 12 structures and 15 rats); right: same but from early to late sleep deprivation. **c,** number and total cumulative duration of sleep intrusions during sleep deprivation (n=20 probes from 15 rats). **d,** distribution of the duration of sleep intrusions in the last hour of sleep deprivation. **e,** further increase in OFF occurrence across sleep deprivation (above and beyond the increase illustrated in b, right) after including sleep intrusions. **f,** change in delta across the experimental conditions indicated in a. In f-h, n=29 unique subject-structure pairs from 12 structures and 15 rats. Bars above plots indicate post hoc general linear hypothesis tests performed on mixed-effects models where asymptotic likelihood ratio tests found a significant (p < .05) omnibus effect of condition. All tests were two-sided and corrected for multiple comparisons (see Methods). Gray bars indicate post hoc tests performed but not significant. Black bars indicate significant post hoc tests (*: p < .05; **: p < .01; ***: p < .001). Models were of the form y ∼ condition + (1 | subject) + (1 | structure) + (1 | subject:structure), with one observation per structure per rat in each condition, so that rat, structure and rat-structure combination each had its own intercept. Data are shown with random effects removed. Precise p-values, confidence intervals, estimates, effect sizes, and model details for f-h are provided in Supplementary Tables S0–S1. **g**, change in temporal and spatial features of all OFF periods across experimental conditions. “Total area norm.” is the total OFF area across the entire condition, normalized by each structure’s maximum possible span and the total condition duration, providing a measure of “total OFFness” that can be compared across subjects and structures. **h,** Changes in small OFF and medium+large OFF rates. Whereas changes in most temporal and spatial features of OFF periods across experimental conditions are qualitatively similar for small OFFs and medium+large OFFs (see Supplementary Fig. 5 for results faceted by small and medium+large OFFs separately), changes in overall OFF differ by size class: during periods of reduced sleep pressure there are fewer medium+large OFF periods, but more small OFF periods.

**Figure 3.**
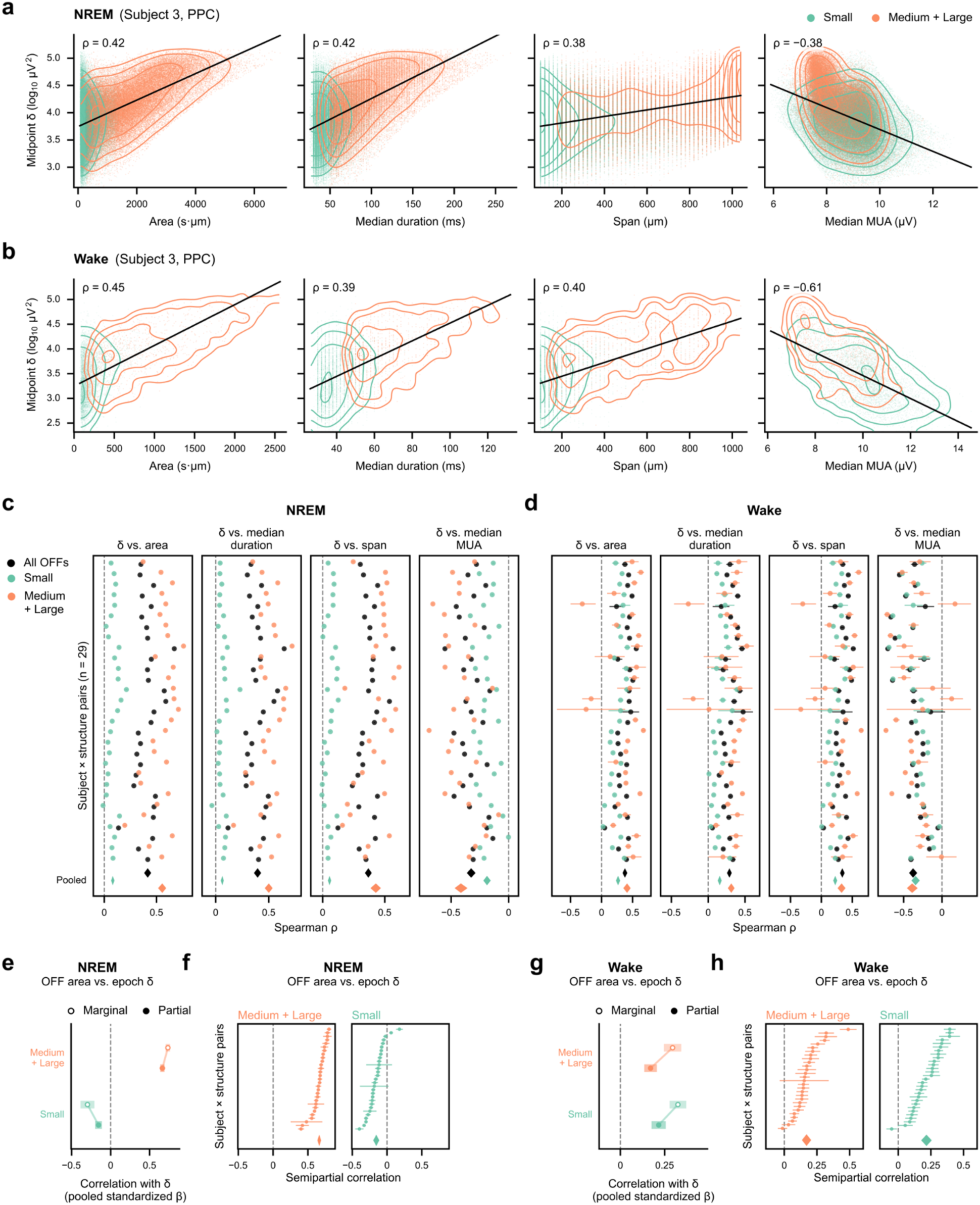
Small OFF periods’ contribution to the prediction of delta activity (δ). **a,** density plots and correlation of per-OFF instantaneous delta power at the OFF period’s midpoint (see Methods) with per-OFF area, median duration, span, and median MUA, in one example cortical region during NREM sleep (Subject 3, PPC). Spearman rho is shown in the upper left of each plot. **b,** same for wake, where OFF periods surviving the strictest selection criteria are uncommon. **c-d,** group-level summary of Spearman correlation analyses illustrated in (a) and (b), respectively (Supplementary Table S2a). Each coefficient is computed within one subject-structure pair (n=29 unique subject-structure pairs from 12 structures and 15 rats). Error bars represent 95% confidence intervals. Diamonds represent the pooled rho for each OFF class (all, small, or medium/large), and their width represents their 95% confidence interval. Pooling is a random-effects meta-analysis across pairs, with no separate term for rat identity (see Methods). **e,** standardized coefficients from linear models predicting the mean delta power of each 10-s epoch from the total OFF area in that epoch (see Methods). Open markers, marginal coefficients from a model with that size class alone; filled markers, semipartial coefficients, which measure what that class adds once the other is accounted for. Coefficients are pooled across pairs by the same random-effects meta-analysis as in (c-d) (n=28-29 pairs; Supplementary Table S2b), and bars are 95% confidence intervals. Nonzero semipartial coefficients indicate that OFF periods in that class add predictive value; the sign of the coefficient is the direction of the relationship between delta and OFF area. **f,** subject-and-structure-level semipartial coefficients underlying the group-level data in (e), for medium/large (left) and small (right) OFF periods. Rows are sorted independently in each plot. The diamond indicates the random-effect pooled estimate, and its width the 95% confidence interval. **g-h,** same as (e-f), for wake.

For analyses that compare delta power across structures and subjects, the per-structure power time series was log-transformed and z-scored relative to the distribution of all artifact-free samples across the entire 48-hour recording. To annotate an OFF period with a co-occurring delta value, we took the log-transformed instantaneous delta power at the sample nearest the event’s midpoint. We used the midpoint value because, unlike the total or the maximum over the event, it does not grow with the number of samples the event spans.

For the epoched partial-correlation analyses (below), NREM sleep and wake were each divided into non-overlapping 10-s epochs; an epoch was retained only if at least 80% of its samples were artifact-free and scored as that state. The delta value of an epoch was the mean of the log-transformed instantaneous delta power over its samples of that state.

### Sleep intrusion scoring

Sleep was scored throughout the recording as described in [10], using LFPs and a synthetic electromyogram (EMG). During sleep deprivation, animals occasionally entered brief episodes of sleep (‘sleep intrusions’) despite behavioral wakefulness (for example, open eyes and upright posture), particularly in the final hour. So that OFF periods in unambiguous wake could be distinguished from OFF periods within sleep intrusions, which were analyzed separately (Fig. 2c-e), the 6 hours of enforced waking were scored a second time, at higher scrutiny, by a second scorer who also had access to sorted unit activity from every recorded brain area (including thalamus, striatum, etc.). For all primary analyses the two wake conditions comprised only epochs scored as wake, so that intrusions were excluded. To assess the contribution of sleep intrusions to the wake OFF-period increase (Fig. 2e), we performed a sensitivity analysis in which OFF periods occurring within intrusions (using NREM detection thresholds) were progressively re-admitted as a function of a maximum intrusion-duration threshold, swept from 1 to 20 s.

### Definition of statistical conditions

To assess the homeostatic regulation of OFF periods and cortical delta activity, we compared six experimental conditions, each defined relative to the 48-hour protocol (Fig. 2a). Early and late sleep deprivation were defined as the first and last cumulative hours of the 6 h sleep deprivation period. Early recovery sleep was defined as the first cumulative hour of NREM sleep following the end of sleep deprivation, and late recovery sleep as the final cumulative hour of NREM sleep before lights-off (9 p.m.). Early baseline sleep was defined as the first cumulative hour of NREM sleep following baseline lights-on. Late baseline sleep (which we also call the circadian-matched baseline) was defined as all NREM sleep on the baseline day within the clock-time window spanned by early recovery sleep 24 h later, widened by 30 min at each end so that enough baseline NREM sleep was available even when the strict window contained little. Unlike the other five conditions, it was therefore not exactly one hour long (median 80 min, range 51–93 min).

### Statistical analysis

Unless otherwise noted, group-level effects were assessed with linear mixed-effects (LME) models [14], in which a response variable was related to a fixed effect of interest while random effects modeled the non-independence of observations from the same subject and the same cortical structure. All LME models were fit by numerical maximum likelihood with the lme4 package [15] in R. The significance of a fixed effect was assessed by an asymptotic likelihood ratio test against the same model with that effect removed. Effect sizes for these tests were quantified as Cohen’s *f*², computed from the change in *R*² between the full and reduced models; because *R*² is not defined straightforwardly for LME models, we followed Selya et al. [16], subtracting the estimated random effects from the response and refitting ordinary least-squares models with and without the fixed effect to obtain comparable *R*² values. Where a significant omnibus effect was found, post hoc comparisons were performed with the multcomp package [17], using the single-step (multivariate-*t*) method to correct for multiple comparisons within each family of contrasts; all tests were two-sided. Post hoc effect sizes were reported as a Cohen’s-*d* analogue, the contrast estimate divided by the square root of the summed variance components of the model. Residual plots of every model were inspected for departures from constant variance and normality. Where these were evident, the response was transformed (log or square root) and/or the model was fit with inverse-variance weights, each observation weighted by the reciprocal of the sample variance of the (transformed) response within its condition; the transform and the weights were chosen per response variable on the basis of these diagnostics. Details of all models and tests are included in Supplementary Tables S0–S5. All original code needed to reproduce our results and figures is available at https://github.com/CSC-UW/cnpix-local-sleep.

Homeostatic changes in delta power and in the properties of OFF periods across the six conditions were assessed with an LME model of the form *y* ∼ condition + (1 | subject) + (1 | structure) + (1 | subject:structure), in which condition was a categorical fixed effect and the crossed random effects allowed each subject, each structure and each subject–structure combination its own intercept (equivalently, structure was treated as a random effect interacting with subject, rather than as a fixed effect). Because OFF period properties exhibit strong temporal autocorrelation, each response variable was reduced to a single value per subject, structure and condition (e.g., median per-OFF area across all early recovery NREM OFF periods detected in Subject 3’s posterior parietal cortex). The omnibus effect of condition was tested against the intercept-only model and, when significant, five pre-specified contrasts were tested post hoc: early versus late wake during sleep deprivation; early recovery sleep versus its circadian-matched baseline (the NREM rebound); early recovery versus early baseline sleep; early versus late recovery sleep; and early baseline versus circadian-matched baseline sleep (Supplementary Tables S0*–*S1).

OFF-period rate and total normalized OFF area, which are near zero in wake, were modeled separately for NREM sleep and for wake, because the pooled six-condition model violated its assumptions for these two measures (Supplementary Methods, “Separate sleep and wake models for OFF rate and total area”).

### Per-event correlations between OFF properties and instantaneous delta power

To test whether the properties of individual OFF periods track delta power on an event-by-event basis, each OFF period was annotated with its co-occurring delta value (see “Delta power computation” above). Within each subject-structure combination, we computed the Spearman rank correlation between each OFF property (area, median duration, span and median MUA) and this per-event delta value, requiring at least four OFF periods per correlation. Per-combination correlations were combined into a single pooled estimate by a random-effects meta-analysis [19] on the Fisher *z*-transformed coefficients: each correlation was weighted by the inverse of its variance plus the estimated between-combination variance, and the pooled coefficient and its 95% confidence interval were computed analytically on the *z* scale and back-transformed to a correlation. Confidence intervals for individual correlations were the corresponding analytic Fisher-*z* intervals. Multiple comparisons across the four OFF properties were controlled by Bonferroni correction. Results are summarized in Supplementary Table S2a. The same analysis was repeated with instantaneous power in the 2–6 Hz and 5–10 Hz bands, computed as described for delta (see “Delta power computation”) with only the passband changed (Supplementary Table S2c). Robustness to influential structures was verified by leave-one-structure-out re-analysis (not shown). For display only, the plotted scatter and regression lines excluded points above the 99.5th percentile of Cook’s distance within each structure, but all reported correlation coefficients were computed from the full set of OFF periods.

To ask whether each property carries information about delta that the other three do not, we also regressed the per-event delta value on all four properties jointly within each subject-structure combination (ordinary least squares on ranks z-scored within the combination), and computed each property’s semipartial correlation with delta: the correlation of delta with the part of that property not predicted by the other three. Its square is the increment in *R^2^* that the property contributes over the other three, and the variance shared among the properties was taken as the full-model *R^2^*minus the sum of the four increments. Semipartial coefficients were pooled by the same random-effects procedure, applied to the coefficients directly, with each coefficient’s variance taken from the regression (Supplementary Table S2d).

### Epoched partial correlations between OFF area and delta power

To test whether small OFF periods carry information about delta power beyond that carried by larger ones, we related epoch-level delta power to the OFF-period area present in the same epoch, separately in NREM sleep and in wake. For each subject-structure combination and each 10-s epoch (see “Delta power computation”), we summed the area of OFF periods in each size class (small and medium/large) and predicted each epoch’s mean log delta power from these per-epoch total areas using linear regression models. Predictors and response were z-scored within each subject-structure combination and state, so that all coefficients are standardized betas. In a ‘marginal’ model, delta was predicted from the area of a single class alone; because the variables are standardized, this coefficient is a correlation and its square is that class’s *R^2^*. To ask whether one class predicts delta beyond the other, we computed the semipartial (part) correlation: delta power and the class of interest were each residualized on the other class, and the delta residuals regressed on the standardized residuals of that class’s area. The square of this semipartial coefficient is the increment in *R^2^* that class contributes over the other alone, so the marginal and semipartial coefficients are the signed square roots of the marginal and incremental variance explained. Because delta power is autocorrelated on the scale of minutes, standard errors were computed with a heteroskedasticity- and autocorrelation-consistent (Newey–West) estimator [20], with a lag corresponding to ∼5 min. Subject-structure coefficients were pooled by the same random-effects meta-analysis used for the per-event correlations, yielding pooled standardized betas and analytic 95% confidence intervals. Results are summarized in Supplementary Table S2b. The analysis was repeated with rank-transformed variables, and with epoch durations of 4 and 30 s in NREM sleep and of 4, 30, 60, and 120 s in wake; coefficients kept their sign and significance in every variant.

### OFF locality/overlap

To quantify the extent to which an OFF period in one cortical structure coincides with OFF periods elsewhere, we defined, for each OFF period, its overlap degree: the number of other simultaneously recorded structures that had at least one temporally overlapping OFF period. An OFF period was termed ‘local’ if its overlap degree was 0 and ‘overlapping’ if it was at least 1. Because longer and more frequent OFF periods overlap by chance more often, we compared the observed overlap against a null model that preserves each structure’s OFF statistics while destroying genuine cross-structure synchrony. In this windowed-shift null, the onset of every OFF period in each partner structure was independently displaced by an amount drawn uniformly from plus or minus half of a 60-s window (reflected at the boundaries of its NREM bout, and preserving each event’s duration), while the structure under test was held fixed; this preserves each structure’s OFF count, duration distribution and within-bout density but scrambles fine cross-structure alignment, and was recomputed over 200 shuffles. An independent analytic check, in which each partner structure is modeled as a homogeneous Poisson process of OFF periods (Fig. 4c), is described in the Supplementary Methods. For each OFF period, its excess overlap was the observed overlap degree minus the mean degree over the windowed-shift shuffles.

**Figure 4.**
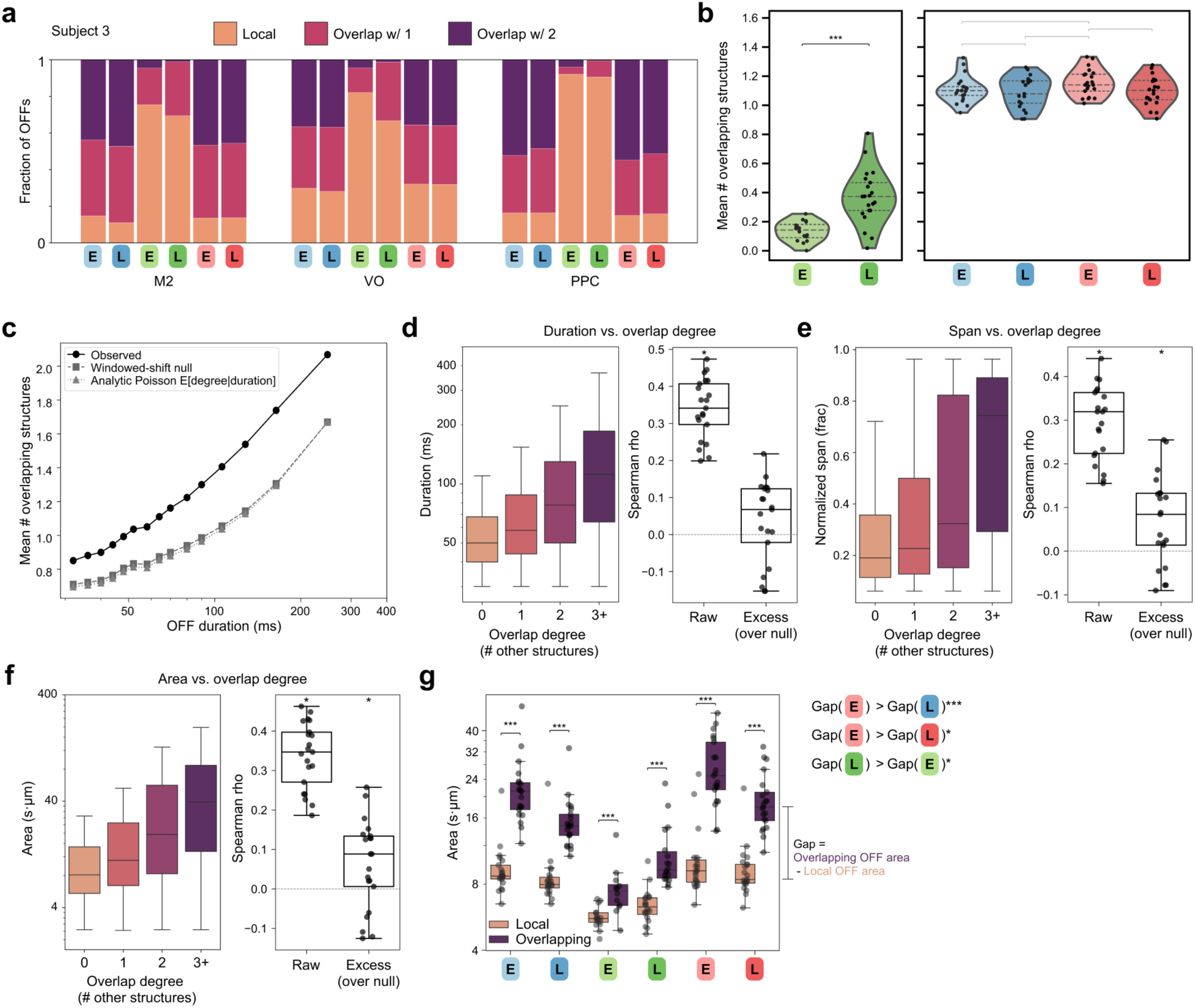
The degree of OFF overlap increases with sleep deprivation and predicts OFF size. **a**, the fraction of OFF periods in a given structure that overlap with an OFF period in exactly 0 (“local”), 1, or 2 other structures in one representative subject. **b,** group-level (n=21 unique subject-structure pairs from 7 unique subjects and 12 unique structures; subjects without multiple structures were excluded from this analysis) summary of OFF locality by condition (Supplementary Table S3a). Bars above plots indicate post hoc general linear hypothesis tests performed on mixed-effects models, conditional on a significant omnibus effect of condition (as assessed by asymptotic likelihood ratio tests). All tests are two-sided and corrected for multiple comparisons (see Methods). Gray bars indicate post hoc tests performed but not significant. Black bars indicate significant comparisons. The model was of the form y ∼ condition + (1 | subject) + (1 | structure) + (1 | subject:structure), with one observation (the mean overlap degree) per structure per rat in each condition, so that rat, structure and rat-structure combination each had its own intercept. Data are shown with random effects removed. **c,** compared to a windowed-shift null that preserves each structure’s OFF count, durations, and within-bout density while destroying fine cross-structure temporal alignment (squares), or to a model in which each partner structure is an independent homogeneous Poisson OFF process with an empirically derived event rate and mean duration (triangles), observed OFF periods exhibit a greater degree of overlap with other structures than expected (Supplementary Table S3a). Observed and null overlap degree were compared within each subject-structure pair by a mixed-effects model with random intercepts for rat and for the rat-structure pair (y ∼ observed vs. null + (1 | subject) + (1 | subject:structure)). **d,** more global OFF periods are associated with longer OFF durations (left), and this relationship can be quantified as the Spearman correlation between each OFF’s duration and overlap degree (right plot, “raw”). “Excess” correlation is the Spearman correlation between each OFF’s duration and its excess overlap, i.e. its observed overlap degree minus the mean degree expected under the windowed-shift null from (c). Each correlation is computed within one subject-structure pair and pooled across pairs (k=21) by a random-effects meta-analysis, with no separate term for rat identity (see Methods). **e,** same as (d), but for normalized span (the span divided by each structure’s maximum possible span). Note that excess correlation is lower than raw correlation, even though increasing span per se does not induce mechanical correlation, because span and duration are highly correlated themselves, and the null preserves this contingency. **f,** same as (d) and (e), but for area. Coefficients, confidence intervals, and p-values for (d-f) are provided in Supplementary Table S3b. **g,** within each condition, overlapping OFF periods are associated with greater area than structure-local OFF periods, and the gap between local OFF area and overlapping OFF area widens in higher sleep pressure conditions (Supplementary Table S3a). Bars above plots indicate within-condition simple effects, conditional on a significant interaction between overlap and condition and a significant main effect of condition. Significance for gap differences is assessed by difference-in-differences contrasts from the interaction model. The interaction model was of the form y ∼ condition × overlap status with the same three random intercepts as in (b), fit to one value per structure, rat, condition and overlap status. All tests are corrected for multiple comparisons (see Methods).

To test whether OFF periods overlapped more than expected by chance, observed and null overlap degrees were compared with an LME model of the form value ∼ quantity + (1 | subject) + (1 | subject:structure), where quantity distinguished observed from null values, and the effect of quantity was tested by likelihood ratio test (Supplementary Table S3a). The dependence of OFF size on globality (Fig. 4d–f) was quantified as the Spearman correlation between each OFF-period’s overlap degree and its duration, span or area; the ‘excess’ correlation is the same correlation computed with each OFF period’s excess overlap in place of its overlap degree, which removes the component attributable to duration and rate alone (Supplementary Table S3b). Both were computed within each subject-structure combination and pooled by the same random-effects meta-analysis as the per-event correlations. Finally, within each condition we compared the size of local and overlapping OFF periods with an LME model of the form response ∼ overlap_status + (1 | subject) + (1 | structure) + (1 | subject:structure), and tested whether the difference in size between overlapping and local OFF periods varied across conditions with difference-in-differences contrasts derived from the corresponding condition-by-overlap-status interaction model (Supplementary Table S3a).

### Correlation between sleep deprivation and subsequent NREM rebound

To test whether the OFF periods expressed during sleep deprivation predict the subsequent homeostatic rebound of NREM sleep, we asked, across subject-structure combinations, whether regions that produced more OFF periods during sleep deprivation showed a larger or a smaller rebound in the sleep that followed. For each combination we computed the OFF-period rate, and separately the total normalized OFF area, over the 6-h sleep deprivation window (the sleep deprivation predictor), and the rebound of the same measure, defined as its value in early recovery NREM sleep minus its value in the circadian-matched baseline. Because sleep deprivation contained brief sleep intrusions (see “Sleep intrusion scoring”), we also recomputed the predictor over wake epochs only, as a sensitivity analysis; the two versions gave the same result (Supplementary Table S4a). We then fit an LME model of the form *y* ∼ *x* + (1 | subject) and tested the slope using a likelihood ratio test against the intercept-only model *y* ∼ 1 + (1 | subject). Its 95% confidence interval was a Wald interval, and its effect size was expressed as Cohen’s *f*² (Supplementary Table S4a). Because high-leverage points could have driven the fits, we refit each model after leaving out each subject, each structure and each subject-structure combination in turn, and every subset of two or three subjects, asking in each case whether the slope remained significant, and computed Cook’s-distance influence diagnostics (Supplementary Table S4b); we also refit each model with rank- and log-transformed predictors (Supplementary Table S4a).

## Results

### Morphological detection resolves spatially restricted and irregularly shaped OFF periods

We implemented a new method to detect OFF periods that is based on the envelope of high-frequency (0.3-5 kHz) activity, rather than spike detection (see Methods; Supplementary Fig. 1). Unlike traditional methods that classify an entire population of neurons (often layer 5 pyramidal cells, sometimes a whole cortical column) as being ON or OFF together based on their pooled activity, our “morphological detection” approach leverages the dense sampling and high signal-to-noise ratio of Neuropixels probes to capture periods of strongly reduced firing with spatially restricted and/or irregular depth profiles during both sleep and wake (Fig. 1a-e; Supplementary Fig. 1). We also implemented novel, spatially resolved extensions of traditional spike-based OFF detection but selected morphological detection after qualitative and quantitative comparison to manually segmented OFF periods (see Supplementary Methods and Supplementary Tables S6a– c). For each OFF period we calculate several parameters that together capture the temporal and spatial extent of the event (Fig. 1f): median duration (per-OFF median duration across involved channels), span (per-OFF difference between the depths of the deepest and most superficial involved channels), and area (the size of each OFF period’s mask). We also introduce a new parameter “median multiunit activity (MUA)” (per-OFF median MUA trace value across all involved channels and samples) that measures the residual firing activity present during the OFF period (lower MUA = lower activity = a more silent OFF period).

### OFF periods occur during extended wake in every cortical area sampled and increase with time awake

Morphologically detected OFF periods were identified throughout chronic, 48-hour long recordings that included a baseline day followed by 6 hours of sleep deprivation and 18 hours of recovery (Fig. 2a). We used two sets of criteria, relaxed (median duration ≥25 ms, total duration ≥30 ms, minimum span ≥100 µm) or stringent (median duration ≥50 ms, total duration ≥60 ms, minimum span ≥200 µm), to classify OFF events by their temporal and spatial extent (Fig. 1f): small (30–59 ms or 100–199 µm) and medium/large (≥60 ms and ≥200 µm). Across the 11 areas (and claustrum) that we sampled in 15 rats, spanning from frontal to occipital cortex, we found a continuum of OFF sizes that varied in the course of extended wake and sleep. Specifically, in the last as compared to the first hour of sleep deprivation, the total OFF count appears to be higher for both small and medium/large events (Fig. 2b, right). Wake at the same clock times on the baseline day showed no such increase (Supplementary Fig. 3). All analyses of early and late sleep deprivation were done after carefully removing even the briefest “sleep intrusions”: short periods of sleep that sometimes occurred during sleep deprivation – especially the final hour – despite behavioral wakefulness (e.g., open eyes and upright posture) (Fig. 2c,d). Comparing the increase in total count of OFF periods without (Fig. 2b, right) and with intrusions (Fig. 2e) shows that the former, i.e. events occurring during unambiguous wake, are the majority, underscoring that local sleep in wake is not a marginal phenomenon (mean OFF count during late sleep deprivation without intrusions = 339.5; with intrusions = 570.2). The increase in total OFF count in late compared to early sleep deprivation appears to occur in all tested areas, with no obvious differences based on an anatomical gradient (Table 1). Visual cortex does exhibit fewer OFF periods during sleep deprivation (Fig. 2b, right), but this is also the case during sleep (Fig. 2b, left; Table 1). We also asked whether OFF periods preferentially engage particular cortical depths, and found a modest bias toward more superficial channels (Supplementary Fig. 4). In sum, during extended wake the occurrence of OFF periods is a significant and broad phenomenon across the cerebral cortex.

**Table 1.**
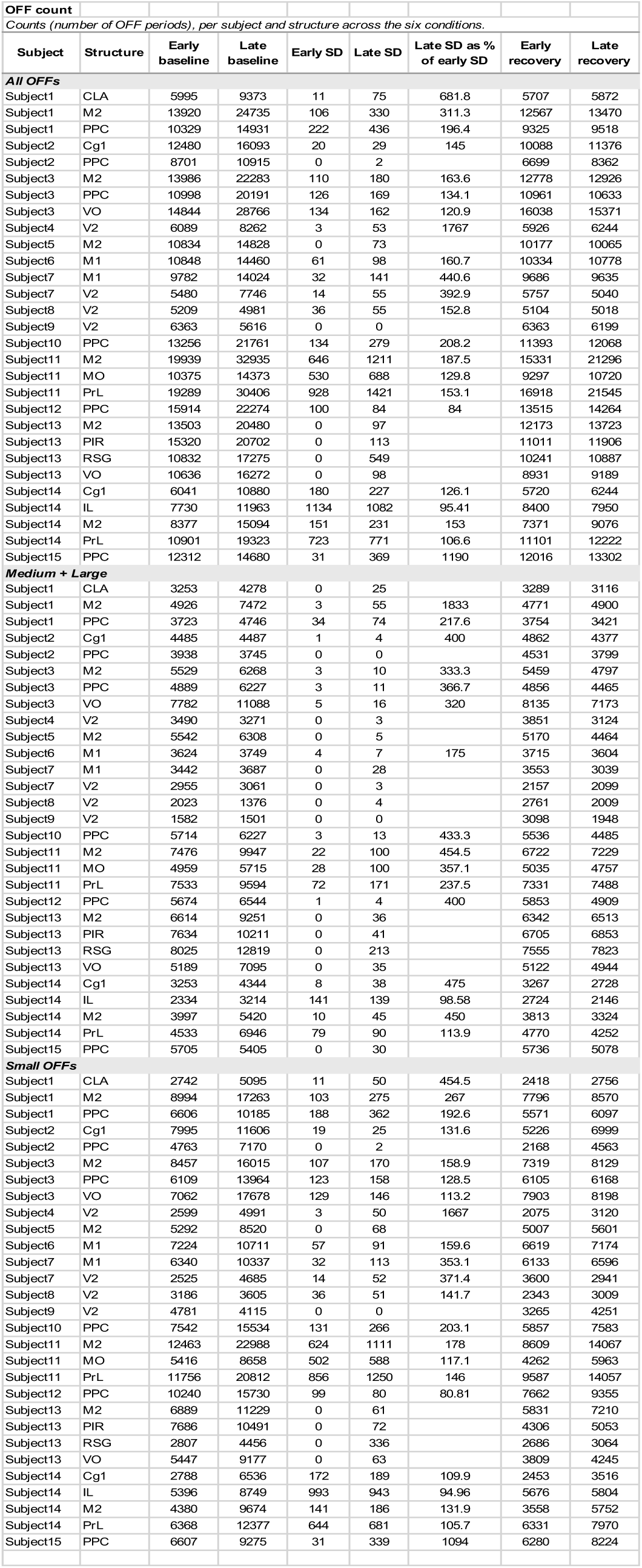
OFF counts per subject and structure across the 6 experimental conditions.

| Subject | Structure | Early baseline | Late baseline | Early SD | Late SD | Late SD as % of early SD | Early recovery | Late recovery |
|---|---|---|---|---|---|---|---|---|
| ***All OFFs*** |  |  |  |  |  |  |  |  |
| Subject1 | CLA | 5995 | 9373 | 11 | 75 | 681.8 | 5707 | 5872 |
| Subject1 | M2 | 13920 | 24735 | 106 | 330 | 311.3 | 12567 | 13470 |
| Subject1 | PPC | 10329 | 14931 | 222 | 436 | 196.4 | 9325 | 9518 |
| Subject2 | Cg1 | 12480 | 16093 | 20 | 29 | 145 | 10088 | 11376 |
| Subject2 | PPC | 8701 | 10915 | 0 | 2 |  | 6699 | 8362 |
| Subject3 | M2 | 13986 | 22283 | 110 | 180 | 163.6 | 12778 | 12926 |
| Subject3 | PPC | 10998 | 20191 | 126 | 169 | 134.1 | 10961 | 10633 |
| Subject3 | VO | 14844 | 28766 | 134 | 162 | 120.9 | 16038 | 15371 |
| Subject4 | V2 | 6089 | 8262 | 3 | 53 | 1767 | 5926 | 6244 |
| Subject5 | M2 | 10834 | 14828 | 0 | 73 |  | 10177 | 10065 |
| Subject6 | M1 | 10848 | 14460 | 61 | 98 | 160.7 | 10334 | 10778 |
| Subject7 | M1 | 9782 | 14024 | 32 | 141 | 440.6 | 9686 | 9635 |
| Subject7 | V2 | 5480 | 7746 | 14 | 55 | 392.9 | 5757 | 5040 |
| Subject8 | V2 | 5209 | 4981 | 36 | 55 | 152.8 | 5104 | 5018 |
| Subject9 | V2 | 6363 | 5616 | 0 | 0 |  | 6363 | 6199 |
| Subject10 | PPC | 13256 | 21761 | 134 | 279 | 208.2 | 11393 | 12068 |
| Subject11 | M2 | 19939 | 32935 | 646 | 1211 | 187.5 | 15331 | 21296 |
| Subject11 | MO | 10375 | 14373 | 530 | 688 | 129.8 | 9297 | 10720 |
| Subject11 | PrL | 19289 | 30406 | 928 | 1421 | 153.1 | 16918 | 21545 |
| Subject12 | PPC | 15914 | 22274 | 100 | 84 | 84 | 13515 | 14264 |
| Subject13 | M2 | 13503 | 20480 | 0 | 97 |  | 12173 | 13723 |
| Subject13 | PIR | 15320 | 20702 | 0 | 113 |  | 11011 | 11906 |
| Subject13 | RSG | 10832 | 17275 | 0 | 549 |  | 10241 | 10887 |
| Subject13 | VO | 10636 | 16272 | 0 | 98 |  | 8931 | 9189 |
| Subject14 | Cg1 | 6041 | 10880 | 180 | 227 | 126.1 | 5720 | 6244 |
| Subject14 | IL | 7730 | 11963 | 1134 | 1082 | 95.41 | 8400 | 7950 |
| Subject14 | M2 | 8377 | 15094 | 151 | 231 | 153 | 7371 | 9076 |
| Subject14 | PrL | 10901 | 19323 | 723 | 771 | 106.6 | 11101 | 12222 |
| Subject15 | PPC | 12312 | 14680 | 31 | 369 | 1190 | 12016 | 13302 |
| ***Medium + Large*** |  |  |  |  |  |  |  |  |
| Subject1 | CLA | 3253 | 4278 | 0 | 25 |  | 3289 | 3116 |
| Subject1 | M2 | 4926 | 7472 | 3 | 55 | 1833 | 4771 | 4900 |
| Subject1 | PPC | 3723 | 4746 | 34 | 74 | 217.6 | 3754 | 3421 |
| Subject2 | Cg1 | 4485 | 4487 | 1 | 4 | 400 | 4862 | 4377 |
| Subject2 | PPC | 3938 | 3745 | 0 | 0 |  | 4531 | 3799 |
| Subject3 | M2 | 5529 | 6268 | 3 | 10 | 333.3 | 5459 | 4797 |
| Subject3 | PPC | 4889 | 6227 | 3 | 11 | 366.7 | 4856 | 4465 |
| Subject3 | VO | 7782 | 11088 | 5 | 16 | 320 | 8135 | 7173 |
| Subject4 | V2 | 3490 | 3271 | 0 | 3 |  | 3851 | 3124 |
| Subject5 | M2 | 5542 | 6308 | 0 | 5 |  | 5170 | 4464 |
| Subject6 | M1 | 3624 | 3749 | 4 | 7 | 175 | 3715 | 3604 |
| Subject7 | M1 | 3442 | 3687 | 0 | 28 |  | 3553 | 3039 |
| Subject7 | V2 | 2955 | 3061 | 0 | 3 |  | 2157 | 2099 |
| Subject8 | V2 | 2023 | 1376 | 0 | 4 |  | 2761 | 2009 |
| Subject9 | V2 | 1582 | 1501 | 0 | 0 |  | 3098 | 1948 |
| Subject10 | PPC | 5714 | 6227 | 3 | 13 | 433.3 | 5536 | 4485 |
| Subject11 | M2 | 7476 | 9947 | 22 | 100 | 454.5 | 6722 | 7229 |
| Subject11 | MO | 4959 | 5715 | 28 | 100 | 357.1 | 5035 | 4757 |
| Subject11 | PrL | 7533 | 9594 | 72 | 171 | 237.5 | 7331 | 7488 |
| Subject12 | PPC | 5674 | 6544 | 1 | 4 | 400 | 5853 | 4909 |
| Subject13 | M2 | 6614 | 9251 | 0 | 36 |  | 6342 | 6513 |
| Subject13 | PIR | 7634 | 10211 | 0 | 41 |  | 6705 | 6853 |
| Subject13 | RSG | 8025 | 12819 | 0 | 213 |  | 7555 | 7823 |
| Subject13 | VO | 5189 | 7095 | 0 | 35 |  | 5122 | 4944 |
| Subject14 | Cg1 | 3253 | 4344 | 8 | 38 | 475 | 3267 | 2728 |
| Subject14 | IL | 2334 | 3214 | 141 | 139 | 98.58 | 2724 | 2146 |
| Subject14 | M2 | 3997 | 5420 | 10 | 45 | 450 | 3813 | 3324 |
| Subject14 | PrL | 4533 | 6946 | 79 | 90 | 113.9 | 4770 | 4252 |
| Subject15 | PPC | 5705 | 5405 | 0 | 30 |  | 5736 | 5078 |
| ***Small OFFs*** |  |  |  |  |  |  |  |  |
| Subject1 | CLA | 2742 | 5095 | 11 | 50 | 454.5 | 2418 | 2756 |
| Subject1 | M2 | 8994 | 17263 | 103 | 275 | 267 | 7796 | 8570 |
| Subject1 | PPC | 6606 | 10185 | 188 | 362 | 192.6 | 5571 | 6097 |
| Subject2 | Cg1 | 7995 | 11606 | 19 | 25 | 131.6 | 5226 | 6999 |
| Subject2 | PPC | 4763 | 7170 | 0 | 2 |  | 2168 | 4563 |
| Subject3 | M2 | 8457 | 16015 | 107 | 170 | 158.9 | 7319 | 8129 |
| Subject3 | PPC | 6109 | 13964 | 123 | 158 | 128.5 | 6105 | 6168 |
| Subject3 | VO | 7062 | 17678 | 129 | 146 | 113.2 | 7903 | 8198 |
| Subject4 | V2 | 2599 | 4991 | 3 | 50 | 1667 | 2075 | 3120 |
| Subject5 | M2 | 5292 | 8520 | 0 | 68 |  | 5007 | 5601 |
| Subject6 | M1 | 7224 | 10711 | 57 | 91 | 159.6 | 6619 | 7174 |
| Subject7 | M1 | 6340 | 10337 | 32 | 113 | 353.1 | 6133 | 6596 |
| Subject7 | V2 | 2525 | 4685 | 14 | 52 | 371.4 | 3600 | 2941 |
| Subject8 | V2 | 3186 | 3605 | 36 | 51 | 141.7 | 2343 | 3009 |
| Subject9 | V2 | 4781 | 4115 | 0 | 0 |  | 3265 | 4251 |
| Subject10 | PPC | 7542 | 15534 | 131 | 266 | 203.1 | 5857 | 7583 |
| Subject11 | M2 | 12463 | 22988 | 624 | 1111 | 178 | 8609 | 14067 |
| Subject11 | MO | 5416 | 8658 | 502 | 588 | 117.1 | 4262 | 5963 |
| Subject11 | PrL | 11756 | 20812 | 856 | 1250 | 146 | 9587 | 14057 |
| Subject12 | PPC | 10240 | 15730 | 99 | 80 | 80.81 | 7662 | 9355 |
| Subject13 | M2 | 6889 | 11229 | 0 | 61 |  | 5831 | 7210 |
| Subject13 | PIR | 7686 | 10491 | 0 | 72 |  | 4306 | 5053 |
| Subject13 | RSG | 2807 | 4456 | 0 | 336 |  | 2686 | 3064 |
| Subject13 | VO | 5447 | 9177 | 0 | 63 |  | 3809 | 4245 |
| Subject14 | Cg1 | 2788 | 6536 | 172 | 189 | 109.9 | 2453 | 3516 |
| Subject14 | IL | 5396 | 8749 | 993 | 943 | 94.96 | 5676 | 5804 |
| Subject14 | M2 | 4380 | 9674 | 141 | 186 | 131.9 | 3558 | 5752 |
| Subject14 | PrL | 6368 | 12377 | 644 | 681 | 105.7 | 6331 | 7970 |
| Subject15 | PPC | 6607 | 9275 | 31 | 339 | 1094 | 6280 | 8224 |

### OFF period number, size and depth of silence change with sleep pressure in wake and sleep

Delta activity reflects homeostatic sleep pressure, being higher in early than late baseline sleep, higher still in recovery sleep, and tending to increase between early and late sleep deprivation [4], as we also find in our recordings (Fig. 2f). Like delta, most features of morphologically detected OFF periods also change with time spent awake and asleep (Fig. 2g; Supplementary Fig. 5). Specifically, over the course of sleep deprivation, OFF periods – whether small or medium/large – become more frequent and involve more channels (small OFF periods also last longer), resulting in a larger per-OFF and cumulative area. Their median MUA also decreases, indicating that they become “deeper” or “more silent”. Following sleep deprivation, in late as compared to early recovery sleep, total OFF counts show a trend toward a slight increase (Fig. 2g), driven by a significant increase in small OFF events and a significant decrease in medium/large OFF events (Fig. 2h). The latter also become less synchronous in late recovery sleep, i.e. the involved neural population enters the OFF period less synchronously (Supplementary Fig. 5c). This pattern is consistent with the fragmentation of fewer large OFF periods into multiple smaller events. Indeed, we found that in late recovery sleep all OFF events (small and medium/large) span fewer channels and medium/large events are also shorter (reduced median duration), resulting in reduced per-OFF area. This reduced per-OFF area more than offsets any possible increase in OFF count, so that the net effect is a reduction in the cumulative OFF area or “total OFFness”. Similar changes in OFF count and size are seen in late compared to early baseline sleep (Fig. 2g).

### OFF period features covary with delta power event by event, in sleep and in wake

We then tested whether the spatial and temporal features of OFF periods correlate with instantaneous delta power on a per-OFF basis. We found that this was the case for each of the features, for both NREM sleep (Fig. 3a,c) and wake (Fig. 3b,d) across the full 48 hours, and that the degree of correlation was comparable in both states. This was also the case when the two groups of OFF periods identified using more or less strict criteria were tested separately (Fig. 3c,d; Supplementary Table S2a), but for small OFF periods in NREM sleep the correlations with area, median duration and span were weak (pooled rho = 0.06–0.08), and only median MUA was more clearly correlated with delta (pooled rho = −0.19). In wake, small OFF periods were correlated with delta on every feature (pooled rho = 0.16–0.27 for area, median duration and span, and −0.35 for median MUA).

Because local sleep in wake has also been inferred from LFP and EEG activity in other frequency ranges (2–6 Hz in rats [4]; 2–10 Hz [6] or 5–8 Hz [5, 7] in humans), and our OFF periods were detected from multiunit activity alone, we repeated this analysis with power in the 2–6 Hz and 5–10 Hz bands. Across all OFF periods, correlations with 2–6 Hz power were nearly identical to those with delta power in both states (pooled rho within 0.04 of that for delta), and correlations with 5–10 Hz power were similar in wake (within 0.07) but weaker in NREM sleep (by 0.08–0.26; Supplementary Table S2c).

Because the four features are themselves correlated, these marginal correlations do not show that each carries independent information about delta. We therefore entered all four into a rank-based multiple regression within each subject-structure pair. Together they explained about a quarter of the rank variance of delta (mean *R^2^* = 0.25 in NREM sleep and 0.26 in wake), most of it shared among the features (0.18 and 0.15). Median duration, span and median MUA each retained a unique contribution in both states, whereas area retained essentially none once the other three were accounted for (|semipartial r| ≤ 0.02; Supplementary Table S2d).

### Small OFF periods carry information about delta power beyond that in medium/large OFF periods

Summed over 10-s epochs, OFF area also tracked delta power: in NREM sleep, the total area of medium/large OFF periods in an epoch was strongly correlated with the epoch’s mean delta power (r = 0.73), and that of small OFF periods was negatively correlated with it (r = −0.30); in wake, the total area of both groups was positively correlated with delta (r = 0.29 for medium/large and 0.32 for small OFF periods; Fig. 3e,g, open markers; Supplementary Table S2b). Moreover, semipartial correlation analyses found that small OFF periods carry unique information: capturing small OFF events improves prediction of delta activity above and beyond what is possible with medium/large OFF periods only, in both NREM sleep (Fig. 3e,f) and wake (Fig. 3g,h). Of note, however, the sign of the correlation changes: in NREM sleep epochs, once the medium/large OFF area is held fixed, additional small OFF area is associated with *reduced* delta (Fig. 3e,f), whereas in wake it is associated with increased delta (Fig. 3g,h). This is again consistent with small OFF periods in NREM sleep reflecting the fragmentation of their medium/large counterparts, i.e., coming at their expense. In wake, there is no such opportunity cost: the alternative to small OFF periods is generally no OFF periods at all.

### OFF periods become less local with time awake, and larger OFF periods overlap more than rate and duration predict

Next, we quantified the extent to which the presence of an OFF period in one area is associated with an OFF period somewhere else. During sleep deprivation such overlap was rare, but it increased from the first to the last hour of sleep deprivation (Fig. 4b; Supplementary Table S3a). In NREM sleep most OFF periods coincided with an OFF period in at least one other area (Fig. 4a,b), and overlap did not differ significantly among the sleep conditions. In NREM sleep, longer OFF periods were more likely to overlap with OFF periods in other structures, beyond what was expected due to their duration and rate alone (Fig. 4c) and, vice versa, more global OFF periods were associated with longer OFF durations (Fig. 4d). The same positive correlation with OFF globality was found for OFF area and span (Fig. 4e,f). Within each sleep and wake condition, overlapping OFF periods were associated with greater per-OFF area than structure-local OFF periods, and the size of this disparity reflected sleep need (Fig. 4g). Overlapping OFF periods remained larger in span and area than local OFF periods when compared within deciles of OFF duration, although by less. At matched duration, this gap was similar across conditions within each state (Supplementary Fig. 6), so its dependence on sleep need runs through OFF duration, in line with the link between duration and overlap (Fig. 4c,d).

### Regions with more medium/large OFF periods during sleep deprivation show a smaller OFF rebound in recovery sleep

Finally, we asked whether the OFF periods that occur during sleep deprivation help to discharge sleep pressure or merely signal its accumulation. This is difficult to assess without a within-subject control, because it is reasonable to expect a positive relationship between OFF period expression during sleep deprivation and early recovery sleep if both reflect accumulated sleep need, even if waking OFF periods do relieve sleep pressure. We therefore focused on the relationship between OFF period expression during sleep deprivation and OFF period *rebound* during early recovery sleep (the difference from circadian-matched baseline), since the latter is at least partially adjusted for each subject-structure combination’s baseline OFF expression, and more directly reflects the magnitude of deviation from the homeostatic set point. We conjectured that if OFF periods during sleep deprivation only index accumulated sleep need, the tissues generating the largest number of events should show the largest OFF-period rebound in subsequent sleep—a positive relationship; if instead they relieve part of that pressure, the relationship should be weakened, abolished, or reversed. For each cortical region we therefore related OFF-period activity during the 6 hours of sleep deprivation to the homeostatic rebound of the same parameter in early recovery NREM sleep.

We found no positive relationship. In fact, for medium and large OFF periods, the rate of OFF periods during sleep deprivation was negatively correlated with the subsequent rate rebound (Fig. 5b and Supplementary Table S4a): the more such OFF periods a region produced during extended wake, the smaller its rebound in early recovery sleep. We checked whether this relationship was driven by outliers. It was present with or without cortical structure as an additional random effect, preserved under rank- and log-transformation of the predictor, present when the rebound was expressed as a ratio to baseline rather than a difference (Supplementary Table S4c), and retained when OFF periods were restricted to unambiguous wake by excluding brief sleep intrusions. No single animal accounted for it, although it weakened when the two animals with the most OFF periods during sleep deprivation were dropped together (it survived dropping any of the other 104 pairs of animals; Supplementary Table S4b).

The relationship also remained significant when we adjusted, one covariate at a time, for other measures of sleep pressure (the delta-power rebound and delta power in late sleep deprivation), for the fraction of sleep deprivation spent in sleep intrusions, for the region’s anterior-posterior position, and for the detector’s event recall and precision against manual annotation in that region (Supplementary Table S6a; Supplementary Methods; Supplementary Table S4c). Only adjustment for the region’s baseline OFF rate weakened the relationship so that it was no longer significant (Supplementary Table S4c). The baseline rate was positively correlated with the OFF rate during sleep deprivation and negatively correlated with the rebound. Because the rebound is already measured from baseline, this adjustment asks a different question: whether, among regions with similar baseline rates, those with more OFF periods during sleep deprivation had lower OFF rates in early recovery sleep. The other relationships we examined pointed the same way, but less robustly. In every model without covariates the slope was negative, whether OFF activity during sleep deprivation was measured as rate or as total area, in medium and large OFF periods or in all of them, and whether the rebound was that of the same measure or of delta power (Supplementary Tables S4a and S4c). For medium and large OFF periods, total area was not significantly related to its own rebound (Fig. 5d) unless the region’s baseline OFF area was included as a covariate, but it did predict a smaller rebound of delta power, for which OFF rate showed a trend (Supplementary Table S4c). With all OFF periods pooled, including the smallest, the relationships for rate (Fig. 5a) and for total area (Fig. 5c) were each significant in some analyses but not across transformations of the predictor (Supplementary Table S4a).

**Figure 5.**
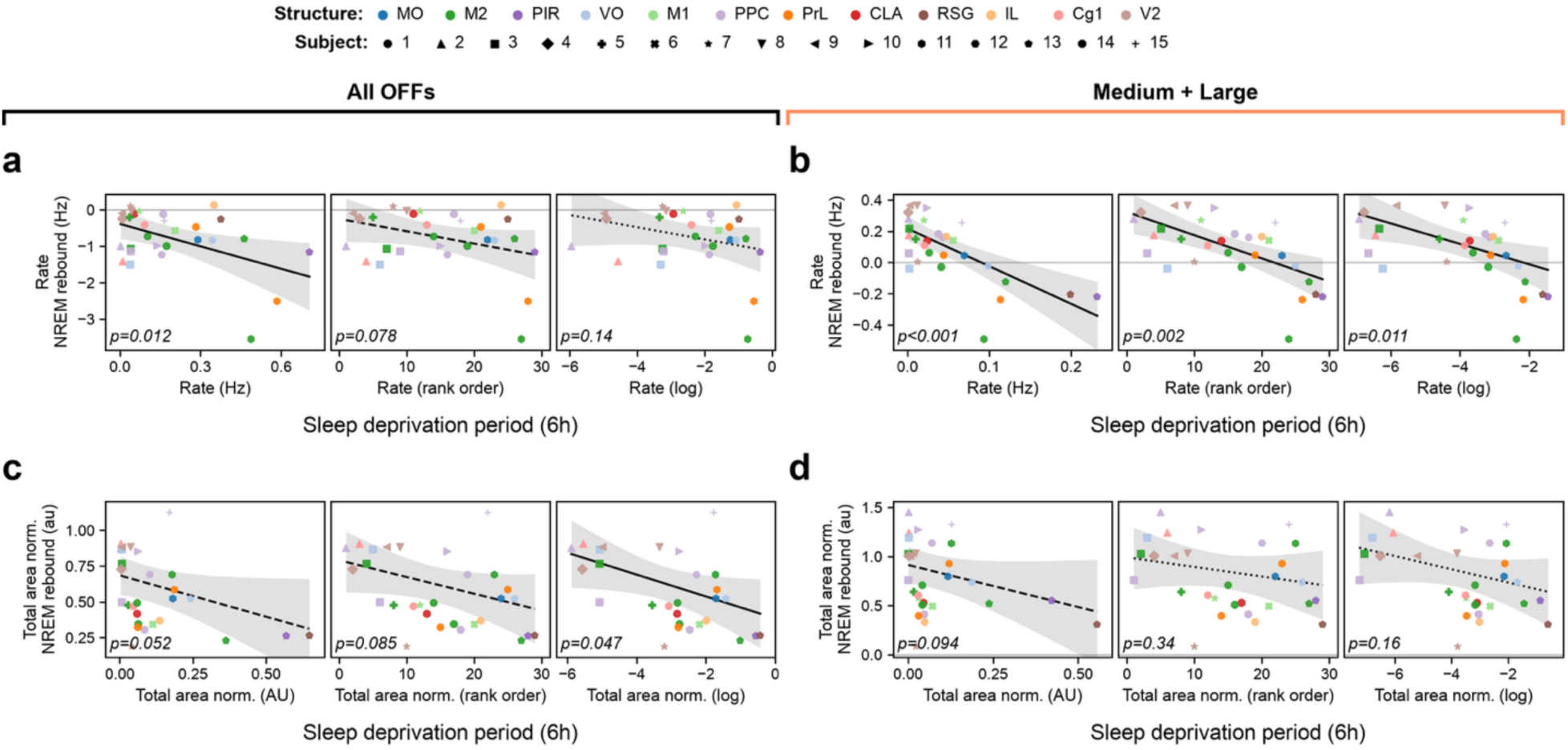
OFF-period activity during sleep deprivation and the subsequent NREM rebound. **a-d,** relationship between OFF-period rate (a,b) or total normalized area (c,d) during the 6 h sleep-deprivation period (x axis) and the homeostatic rebound of the same parameter in early recovery NREM sleep (y axis; its difference from the circadian-matched baseline, as in Fig. 2a), for all OFF periods (a,c) and for medium and large OFF periods only (b,d). Each point is one cortical region in one rat (n=29 subject-structure pairs from 15 animals); the sleep-deprivation predictor and the rebound are both computed for that same region, so each point is a within-region comparison. Color indicates structure and marker shape indicates subject (legends, top). Within each panel, the three plots show the sleep-deprivation predictor untransformed (raw), rank-transformed, and log-transformed; the response is unchanged, and p values (asymptotic likelihood ratio test) are inset. Solid, dashed and dotted lines of fit indicate significant (p <.05), trending (p < .1), and non-significant slopes, respectively. Linear relationships and 95% confidence intervals (shaded regions) are model expectations from linear mixed-effects models with a by-subject random intercept (rebound ∼ sleep-deprivation activity + (1 | subject)), in which structure does not appear as a term because the subject-structure pair is the unit of observation, estimated using a marginal effects method (see Methods for details). See Supplementary Table S4a for slopes, effect sizes (Cohen’s f²) and confidence intervals, and for the equivalent models fit with an additional by-structure random intercept and with sleep deprivation restricted to intrusion-free wake.

## Discussion

Local sleep in wake was first documented in rats [4] and later confirmed in humans at the single-neuron level [6] and using scalp/hd-EEG [5, 8]. These studies left open several major questions, including whether this phenomenon is ubiquitous, whether it affects some cortical areas more than others and whether, despite its documented behavioral cost, it can provide some benefits to the brain. Here, using high-density Neuropixels recordings in the rat and a novel detection method, we found that all tested cortical regions show OFF events during extended wake, with no evidence for some areas being more resilient than others. We also found that the number and spatiotemporal extent of OFF periods increase during sleep deprivation, and that their features track delta activity event by event, in wake as in sleep, consistent with the hypothesis that they reflect sleep need. Finally, we found that regions that produced more medium/large OFF periods during sleep deprivation showed a smaller rebound in recovery sleep, consistent with a partial discharge of sleep pressure before the animal goes to sleep.

We found evidence for local sleep in wake wherever we looked. We also observed that local OFF periods are more frequent in frontal/prefrontal cortex than occipital cortex, but the same gradient was present in early sleep deprivation, late sleep deprivation, as well as during sleep. Moreover, the relative increase in OFF rate between early and late sleep deprivation did not seem to differ between anterior and posterior areas (Table 1). The consistently higher rate of OFF events in frontal areas, whether in wake or in sleep, possibly reflects intrinsic network properties, functional role, and overall plasticity. Indeed, delta activity is already higher in frontal than in occipital cortex during baseline sleep and shows a larger frontal rebound after sleep deprivation, in both rodents and humans [21, 22], and sleep slow waves disproportionately originate in and propagate from anterior regions [23], where cortical gray-matter volume is also greatest. In sum, these results suggest that local sleep in wake is a common consequence of prolonged wake across the cortical mantle and provide no evidence for some areas being more resilient to this phenomenon. Our coverage however was far from being exhaustive, and it remains possible that some cortical areas are immune to local sleep. In humans, slow waves have been reported in every cortical lobe during resting wake [Yang et al., 2026], though in field-potential recordings from patients with epilepsy in which time awake was not controlled; how these relate to the OFF periods described here remains to be established.

OFF periods in wake also behaved as expected if they reflect sleep need. With time awake they became more frequent, longer, larger and quieter, in parallel with delta activity, and at the level of single events their size and residual firing covaried with delta power, in wake as in NREM sleep, and similarly with power in the 2–6 Hz band in which local sleep was first described in rats [4]. Small OFF periods, which conventional methods are not designed to capture, carried information about delta beyond that in medium/large events. In NREM sleep this information was of opposite sign, more small OFF periods at a fixed medium/large area meaning less delta, consistent with small events arising from the fragmentation of larger ones as sleep pressure declines. In wake, where the alternative to a small OFF period is usually no OFF period at all, both classes tracked delta in the same direction. Finally, in sleep, OFF periods in different areas coincided more often than chance alone predicts, overlapping OFF periods were larger, and in wake overlap increased across sleep deprivation, so that local sleep became less local as time awake accumulated. These findings are consistent with the local, use-dependent regulation of sleep slow waves [24–26] and of local sleep in wake [5, 8] described previously, although our protocol did not manipulate the use of specific areas.

The present results also suggest that, in addition to signaling sleep need, OFF periods in wake may contribute to its discharge. Regions that produced more medium/large OFF periods during the 6 h of sleep deprivation showed a smaller rebound of OFF rate in subsequent recovery sleep. The relationship was clearest for medium/large OFF periods and weaker and less robust when small events were included, consistent with the idea that, as long as they are spatiotemporally consolidated, the sporadic local OFF periods of extended wake discharge part of the sleep pressure that would otherwise be dissipated in subsequent NREM sleep, even if their rate is low; the small and more numerous OFF periods may reflect the fragmentation of wake as much as consolidated local sleep. Regions with more total OFF area during sleep deprivation also showed a smaller rebound of delta power (Supplementary Table S4c). These relationships are correlational, however, and do not establish such a discharge. When each region’s baseline OFF rate was added as a covariate it weakened and was no longer significant (Supplementary Table S4c), so part of it may reflect a regional trait, regions that generate many OFF periods in general showing smaller rebounds; if this trait is the same propensity that produces OFF periods in wake, the two readings cannot be separated without manipulating OFF periods directly. Nor can our analysis distinguish whether a discharge would depend on the number of OFF periods or on cumulative OFF activity: across regions, OFF rate and total OFF area during sleep deprivation were nearly collinear, and total area also predicted the rebound of OFF rate (Supplementary Table S4c).

A causal contribution is supported more directly by a recent study in which we optogenetically induced OFF periods for 30 minutes in a small cortical area while previously sleep deprived mice remained awake [27]. This manipulation locally reduced sleep need, as reflected by a decrease in delta activity during subsequent recovery sleep, and provided the affected cortical network with key benefits normally observed after several hours of sleep, including synaptic renormalization and memory consolidation in a task known to depend on cortical slow waves [27]. Whether the OFF periods that occur spontaneously during extended wake afford the same synaptic- and circuit-level benefits is an open question. Their rate is much lower than during optogenetically enforced local sleep, and may be too low to provide substantial long-term benefits while high enough to occasionally impair performance [4–8]. In a mouse model of stroke, optogenetically evoked slow waves improved motor recovery and drove axonal sprouting when induced during sleep, but not when the same number of stimuli was delivered at night, when the animals were mostly awake [28], although OFF induction in wake may have been less effective than in sleep.

The mechanisms underlying local OFF periods in wake remain unclear. Two candidates are disfacilitation and inhibition. Local disfacilitation could stem from declining levels of arousal-promoting neuromodulators or from “fatigue” of cortical pyramidal neurons, but cortical electrical or optogenetic stimulation can trigger short OFF periods in wake [29], which is difficult to explain if reduced synaptic drive were the primary trigger. Cortical fast-spiking (parvalbumin-positive) interneurons are more active in sleep than in wake [30–32], but we are not aware of evidence that they become progressively more active during extended wake, and their peak activity occurs in REM sleep, when slow waves are rare [30–32]. A third possibility is that OFF periods in wake reflect increased neuronal activity and excitability, as in NREM sleep, where sustained synchronous recruitment of cortical neurons during the ON period triggers the subsequent OFF period. Wake-dependent synaptic activity and potentiation would lead to stronger and more synchronous firing of pyramidal neurons and to the progressive recruitment of Martinotti cells, which inhibit all cortical cell types, as seen during OFF periods [27, 33]; rising intracellular chloride during sleep deprivation, as reported in somatosensory cortex ([34] but see [35]), could contribute by enhancing pyramidal excitability and synaptic potentiation [36]. In this view, local OFF periods in wake resemble the isolated slow waves and OFF periods described in awake humans with focal epilepsy, which both slowed cognitive processing and reduced cortical excitability [37]. Distinguishing these possibilities will require causal tests, such as silencing specific interneuron classes during extended wake.

This study has several limitations. First, as mentioned, our sampling was far from exhaustive. Each Neuropixels probe interrogates only a thin column of tissue, and we have yet to analyze subcortical structures such as the thalamus. Our recordings were also restricted to adult male Sprague Dawley rats studied under a single sleep deprivation protocol (6 h), so the extent to which these findings generalize remains to be established. Our detection method also differs from those used in most previous studies: the multiunit envelope is a proxy for spiking, our OFF periods are defined by strongly reduced, not necessarily absent, firing (as captured by the residual median MUA), and the thresholds and size criteria (relaxed versus stringent) are pragmatic, so absolute OFF rates and sizes are not directly comparable across the literature. Moreover, the evidence that OFF periods in wake can discharge sleep pressure is correlative, and both the behavioral costs and the putative restorative benefits of local sleep are inferred from prior work. For the same reason, our data cannot adjudicate the underlying cellular mechanism. These are all open questions that will require targeted, cell-type-specific and causal experiments.

## Acknowledgements.

Supported by NIH grants R01NS131389 (CC, GT), U.S. Department of Defense grant PR230899 (CC), and U.S. Department of Defense grant W911NF1910280 (CC, GT). Generative AI was used to write code, to draft certain figure captions and passages of the Methods, and to draft individual sentences of the Results, all of which were reviewed and edited by the authors.

## Author contributions

GF, MLC collected the data; GF, TB analyzed the data; GF, WM performed statistical analysis; GF, CC, GT designed the experiments; GF, CC and GT wrote the paper.

## Data Availability

The data underlying this article will be shared on reasonable request to the corresponding author.

## Supplementary Methods

### Block-wise threshold estimation

Block-wise thresholding (Methods, “Morphological OFF period detection”) was motivated by the observation that fixed, whole-recording quantiles were unstable. For example, in one representative subject, a threshold pinned to the whole-recording NREM Q20 (20^th^ percentile), regardless of channel depth, swept between Q5 and Q36 of the local (block-level) NREM distribution across blocks, a drift of 31 percentile-equivalent units. Without block-wise thresholding the median Q20 drift across all channels of the 29 subject-structure combinations (15 subjects) was 28 percentile-equivalent units. Not all of this drift is a change in signal scale. When each channel’s threshold was held at its value in one condition, the fraction of samples below it was higher in the condition with greater expected sleep need, by a median factor of 1.35–1.52 in four of the five comparisons tested (Supplementary Fig. 7). Block-wise thresholding removes these changes along with any drift in signal scale.

### Spatially tiled unit-based OFF period detection

As a supplemental, spike-based comparison with morphological detection, we implemented a pair of detectors that operate on sorted-unit spike trains rather than on the MUA envelope. In the “fixed-tiled” variant, to retain spatial resolution while pooling enough spikes for stable detection, we tiled each cortical structure into overlapping depth bands (250-µm-wide, 125-µm step) and, within each band, pooled the spike trains of all units whose depth fell in the band into a single multiunit train binned at 10 ms. OFF periods were detected within each band with a two-state (ON/OFF) Poisson hidden Markov model (HMM) with a ‘sticky’ self-transition constraint enforcing a minimum expected OFF dwell time of 50 ms, in which the OFF state corresponds to near-complete population silence ([12]; ‘on_off_detection’ package; https://github.com/CSC-UW/on_off_detection). Band-level OFF periods were then merged across bands into spatially extended events: proceeding from the longest events, we greedily absorbed shorter events overlapping in both depth (≥50 µm) and time (≥20 ms), retaining the union of their extents, and discarded events shorter than 80 ms after merging.

In the “greedy-tiled” variant, spatial resolution was sacrificed for more uniform sensitivity. Rather than tiling the structure with fixed-width bands, we grew each band from one end of the structure’s depth extent, adding progressively more units until the band’s pooled firing rate exceeded a target (100 Hz) and its depth span exceeded a floor (150 µm); successive bands then overlapped by 75% of pooled firing rate rather than by a fixed depth. Because every band pools a comparable total firing rate, detection sensitivity is approximately equalized across bands, but the bands are correspondingly coarser, of unequal, firing-rate-dependent width, and span only a single spatial scale. Per-band detection (the sticky Poisson HMM) and the across-band merge were otherwise identical to the fixed-tiled variant.

### Agreement between detectors and manual annotation

In order to compare the performance of different OFF detectors, and to obtain a calibrated sense of how far each automated detector’s labels are from human judgement, all OFF periods during the first 3.1–6.9 min (median 3.4 min) of early recovery NREM sleep were manually annotated in napari ([13]; Wacom PTH-651/K digital tablet and stylus) by tracing their spatial (channel) and temporal extent directly on the MUA heatmap images and stored as instance-segmentation masks. We compared automatically detected and manually segmented OFF periods on the common set of detection channels, using the per-pixel mask of each event on the annotation grid (one pixel = 3.3 ms × 20 µm). Agreement was quantified at the pixel level: sensitivity (recall), precision, F1 score and intersection-over-union (IoU). The spatially tiled unit-based detectors were scored in the same way: their OFF periods were rasterized onto the annotation grid and compared with the same manual labels, on the same channels and with the same metrics. Because the pixel-level metrics are conservative by construction, penalizing every boundary disagreement on every channel of every event, we also computed two event-level metrics: event recall, the fraction of manually segmented OFF periods that share at least one pixel with a detection, and event precision, the fraction of detections that share at least one pixel with a manually segmented OFF period.

Morphological detection recovered the manually annotated OFF periods well, and to a similar degree at both admission criteria (F1 0.65 and 0.64; Supplementary Table S6a). At the stringent criterion it marked about as many OFF periods as the annotators did (81 versus 84 per minute), and at the relaxed criterion about twice as many (168 per minute). The relaxed and stringent thresholds traded sensitivity for precision as intended: relaxing the criterion recovered more of the manually labeled pixels and OFF periods (sensitivity 0.58 versus 0.54; event recall 0.93 versus 0.75) at the cost of more detections that matched no manual label (precision 0.78 versus 0.85; event precision 0.59 versus 0.81), with little net change in overall pixel-level agreement (F1 and IoU). Most of this difference came from small OFF periods (those meeting the relaxed but not the stringent criterion), which made up 77% of the relaxed-criterion detections that matched no manual label; per structure, 38 ± 24% of small OFF periods overlapped a manual label, compared with 81 ± 17% of OFF periods meeting the stringent criterion. In sorted spikes, which the detector does not use, the small OFF periods that matched no manual label were nearly as silent as those that matched one: they contained a median of 4% of the spikes expected on the same channels at random times (2% for matched ones). Neither of the two supplemental, spike-based detectors did as well as morphological detection across the annotated structures. The fixed-tiled detector recovered more of the manually labeled pixels (sensitivity 0.74 versus 0.58) but over-detected, producing the most events (232 versus 168 OFF periods per minute) and the lowest precision (0.44; event precision 0.45), and therefore the lowest F1 (0.54) and IoU (0.39). On the 20 structures for which it could be constructed, the greedy-tiled detector’s F1 was slightly higher than that of the relaxed criterion (0.69 vs. 0.66), but only because it marked more of each annotated OFF period (sensitivity 0.69 vs. 0.59), at lower precision (0.73 vs. 0.79), and it recovered fewer of the manually segmented OFF periods (event recall 0.78 vs. 0.92). It also has two drawbacks that F1 does not reflect: its bands are set by firing rate, so its spans and areas are not comparable across structures, and it could not be constructed for the other 8 structures, whose firing rates were too low.

Two further time periods were annotated in the same way (Supplementary Table S6b): NREM sleep at lower sleep pressure, on the baseline day at the clock time of early recovery sleep (the circadian-matched baseline; 13 structures in 6 rats), and wake across the whole 6-h sleep deprivation (16 structures in 9 rats). Morphological detection had the highest F1 with both sets (0.58 and 0.59 for the relaxed and stringent criteria at baseline, 0.57 and 0.56 in wake) and the fixed-tiled detector the lowest. The greedy-tiled detector was less stable: where it could be constructed, its F1 went from 0.69 in early recovery sleep to 0.55 at baseline and 0.52 in wake (relaxed criterion, on the same structures: 0.66, 0.60 and 0.57), and it recovered far fewer of the annotated OFF periods (event recall 0.53 versus 0.91 at baseline, and 0.60 versus 0.89 in wake). Agreement was lower in wake than in early recovery sleep for every detector. Much of this drop is expected from how rare OFF periods are in wake: precision, and with it F1, falls as OFF periods occupy less of the image, even when a detector’s sensitivity and false-positive rate are unchanged. In both sets the relaxed criterion’s F1 rose with the fraction of annotated pixels across structures (Spearman ρ 0.84 at baseline and 0.79 in wake), whereas sensitivity and false-positive rate did not (|ρ| ≤ 0.42, p ≥ 0.10). To estimate the size of this effect, we recalculated the relaxed criterion’s F1 for each wake structure, keeping the sensitivity and false-positive rate measured in that structure but setting the fraction of annotated pixels to the median for early recovery sleep. The recalculated F1 was 0.71, against 0.66 for early recovery sleep by the same calculation.

The last hour of sleep deprivation was also annotated, image by image, by a different annotator. On the 13 structures (8 rats) where this annotation and the sleep-deprivation annotation of Supplementary Table S6b cover the same images, we scored every image, since both annotators had inspected all of them (Supplementary Table S6c). The two annotations agreed only moderately (F1 0.48), mainly because one marked about twice as many OFF periods as the other (1.0 versus 0.5 per minute). The relaxed criterion found 66% of the denser annotation’s OFF periods, more than the sparser annotation did (38%), and 79% of the sparser annotation’s, about as many as the denser one did (81%). Both spike-based detectors agreed far less well with either annotation than morphological detection did (F1 0.01–0.09, against 0.30–0.34), and over-detected: they marked 10 (greedy-tiled) and 42 (fixed-tiled) OFF periods per minute, where the annotators marked 0.5 and 1.0.

Like detected OFF periods, the wake OFF periods marked by the annotators (1,440 in 16 structures) became more frequent, larger, and quieter over sleep deprivation. From the first half of sleep deprivation to the second, their rate rose by 215% (95% confidence interval 116 to 314; p < 0.001) and their cumulative area by 202% (92 to 312; p = 0.002), and their median MUA fell by 1.9% (0.6 to 3.2; p = 0.015; LME models, likelihood ratio tests). The area of individual OFF periods showed a trend toward an increase between halves (8%; p = 0.081) and increased significantly from the first hour of sleep deprivation to the last annotated hour of wake (40%, 6 to 85; p = 0.029). Event by event, they also tracked delta power (pooled Spearman ρ 0.45 for area, 0.43 for duration, 0.24 for span and −0.55 for median MUA; all p ≤ 0.002 after Bonferroni correction).

### Ordinal and Gaussian-mean models of small OFF-period shape

For small OFF periods, median duration and span take only a few discrete values, because span is quantized to the electrode pitch (20 µm) and duration to the MUA sampling interval (2 ms). The median of these quantities for each subject, structure and condition was therefore coarsely discretized, and the residuals of the linear mixed model were visibly striped. We addressed this in two ways. First, the median summaries were modeled with a cumulative-link (ordinal) mixed model. Second, we used the mean instead of the median of the per-OFF quantity (mean median-duration and mean span), which is effectively continuous and met the assumptions of the Gaussian linear mixed model used for the other measures. The two analyses agreed, and we report the Gaussian-mean analyses in Supplementary Fig. 5b and Supplementary Table S1, for consistency with the other measures.

### Separate sleep and wake models for OFF rate and total area

Where possible, response variables were modeled across all six conditions in a single mixed model. OFF-period rate and total normalized OFF area, however, are near zero in wake, where OFF periods are sparse, so pooling wake and sleep produced a floor effect and heteroscedastic residuals that persisted under log and square-root transformation and inverse-variance weighting. For these two measures we therefore fit NREM sleep and wake as separate models, each with its own variance components, which satisfied the model assumptions.

### Poisson null for cross-structure overlap

As an independent analytic check on the windowed-shift null (Methods, “OFF locality/overlap”), we also modeled each partner structure as a homogeneous Poisson process of OFF periods, under which the probability that a focal OFF period of duration d overlaps a given partner structure is 1 − exp(−λ(d + μ)), where λ and μ are that structure’s empirical OFF rate and mean duration; the expected overlap degree is the sum of these probabilities across partner structures (Fig. 4c, triangles).

### Correcting for mechanical correlation of edge asynchrony and span

Onset and offset asynchrony are computed by taking the median absolute deviation (MAD) of onset/offset times across the channels an OFF period spans. The spatial closing operation applied during OFF period detection (see Methods, “Morphological OFF period detection”) is a form of spatial smoothing, and interacts with the MAD to induce a floor effect (MAD is guaranteed to be 0 for any OFF spanning 120 µm or less) that releases with increasing span, introducing a mechanical correlation between OFF span and edge MAD. We confirmed that this dependence is a property of the detector by generating synthetic OFF periods with a fixed, span-independent latent edge dispersion and propagation slope and passing them through the detector, labeling, and measurement code. This simulation reproduced the observed relationship between MAD and span across the full range (RMS error 0.83 ms).

We therefore report span-adjusted asynchrony obtained by marginal standardization [18]. We first estimated how MAD depends on span, as a single curve (scaled to unit std. dev.) shared across all subject–structure pairs and conditions, scaled by a per-pair amplitude coefficient (in ms per std. dev.). Both the shared curve and the amplitudes were estimated only from variation within each pair–condition combination, so neither could absorb a condition effect. Within each pair, and separately for NREM sleep and wake, each OFF period’s MAD was then regressed on condition and on that curve, and the fitted model was used to predict MAD for every one of that pair’s OFF periods under each condition in turn. Averaging those predictions gives one adjusted mean, in milliseconds, per pair–condition combination. Every adjusted mean is therefore an average over the same OFF periods, and hence over the same distribution of spans, so conditions are compared at matched span rather than at whatever spans they happened to contain. When analyzing wake OFF periods, 25 of 42 pair–condition combinations had fewer than 60 OFF periods, which might have made estimation of the amplitude coefficient unreliable, so these pair–condition combinations were given the amplitude fitted across all wake OFF periods at once. The qualitative results were robust to these choices. For example, removing per-pair amplitudes entirely, or replacing the shared curve with a quadratic or a saturated per-channel-count coding, changed the NREM ON-transition rebound (early recovery minus circadian-matched baseline) by at most 0.03 ms, and did not change wake contrast significance or direction. Our estimated ON-transition rebound (−0.44 ms) was also reproduced by different forms of adjustment: stratified direct standardization (−0.41 ms) and an event-level model fitted within each pair and pooled by random-effects meta-analysis with standard errors clustered on 60-s blocks (−0.43 ms).

Adjusted means were analyzed with the same mixed model, likelihood ratio test, post hoc contrasts and effect sizes as every other response variable, and are reported for the medium/large OFFs only, because the aforementioned floor effect precludes complete and confident analysis of the small OFF periods.

### Robustness of the sleep deprivation–rebound slope to potential confounds

The negative relation between OFF-period rate during sleep deprivation and the rebound of OFF-period rate (Methods, “Correlation between sleep deprivation and subsequent NREM rebound”) could arise from differences between subject-structure combinations that affect both quantities. Differences between animals are absorbed by the random intercept for subject in the model y ∼ x + (1 | subject). To address other candidate confounds, we added each of the following covariates to this model in turn, with x computed over the entire sleep deprivation window. As measures of sleep pressure that do not depend on OFF-period detection, we used the combination’s delta-power rebound (mean log-transformed, z-scored delta power in early recovery NREM sleep minus that in the circadian-matched baseline; see Methods, “Delta power computation”) and its mean delta power in late sleep deprivation. We also included the fraction of the sleep-deprivation window occupied by sleep intrusions. For regional differences we used the anterior-posterior coordinate of the structure, and for differences in baseline OFF expression, the combination’s OFF-period rate in circadian-matched baseline NREM sleep. For differences in detector performance we used the structure’s event recall and event precision against manual annotation of early recovery NREM sleep (Supplementary Table S6a), which reflect missed and spurious OFF periods, respectively. In each case the slope of x was tested with a likelihood ratio test against the same model without x. We refit the unadjusted model with the rebound expressed as the log ratio of the early-recovery to the circadian-matched baseline rate, which normalizes the rebound by each combination’s baseline. To compare OFF rate and total OFF area as predictors, we fit the same model with the total normalized OFF area during sleep deprivation as x and the rebound of OFF rate as y, and with OFF rate and total OFF area in turn as x and the delta-power rebound as y. We also refit the total-area relation (Fig. 5c,d) with the baseline total area as a covariate. Associations among the predictor, the rebound and the candidate confounds were measured with Spearman rank correlations across combinations (Supplementary Table S4c).

### Depth profile of OFF involvement

To test whether OFF periods engage some cortical depths more than others, we selected cortical structures that a probe traversed across the cortical layers over a span of more than 500 µm (identified by manual curation), and, for each structure, counted the number of OFF periods (NREM sleep, relaxed criterion) involving each channel as a function of depth. We compared this empirical depth profile to a null that preserves the size and shape of every OFF period and randomizes only its depth. For each shuffle, every detected OFF period was translated rigidly along the depth axis by an integer number of channels drawn uniformly among the positions that keep it within the detection window. The null therefore assumes that an OFF period’s depth is independent of its size and shape and uniform across depth (and that detection sensitivity is uniform across depths). From 200 such shuffles we computed, for each structure, two departure metrics: the total variation between the observed and mean-null depth profiles (the fraction of OFF mass that must be relocated for the null to match the observation) and a signed superficial-versus-deep asymmetry (the observed-minus-null fraction of OFF mass falling in the superficial half of the span, with positive values indicating an excess toward the cortical surface). Per-structure metrics were summarized across structures with intercept-only LME models of the form metric ∼ 1 + (1 | subject). Group means and their 95% confidence intervals were taken from the fixed intercept, with a second confidence interval obtained by a subject-cluster bootstrap that resampled whole subjects (5,000 iterations) as a check (Supplementary Table S5).

---

**Supplementary Figure 1.**
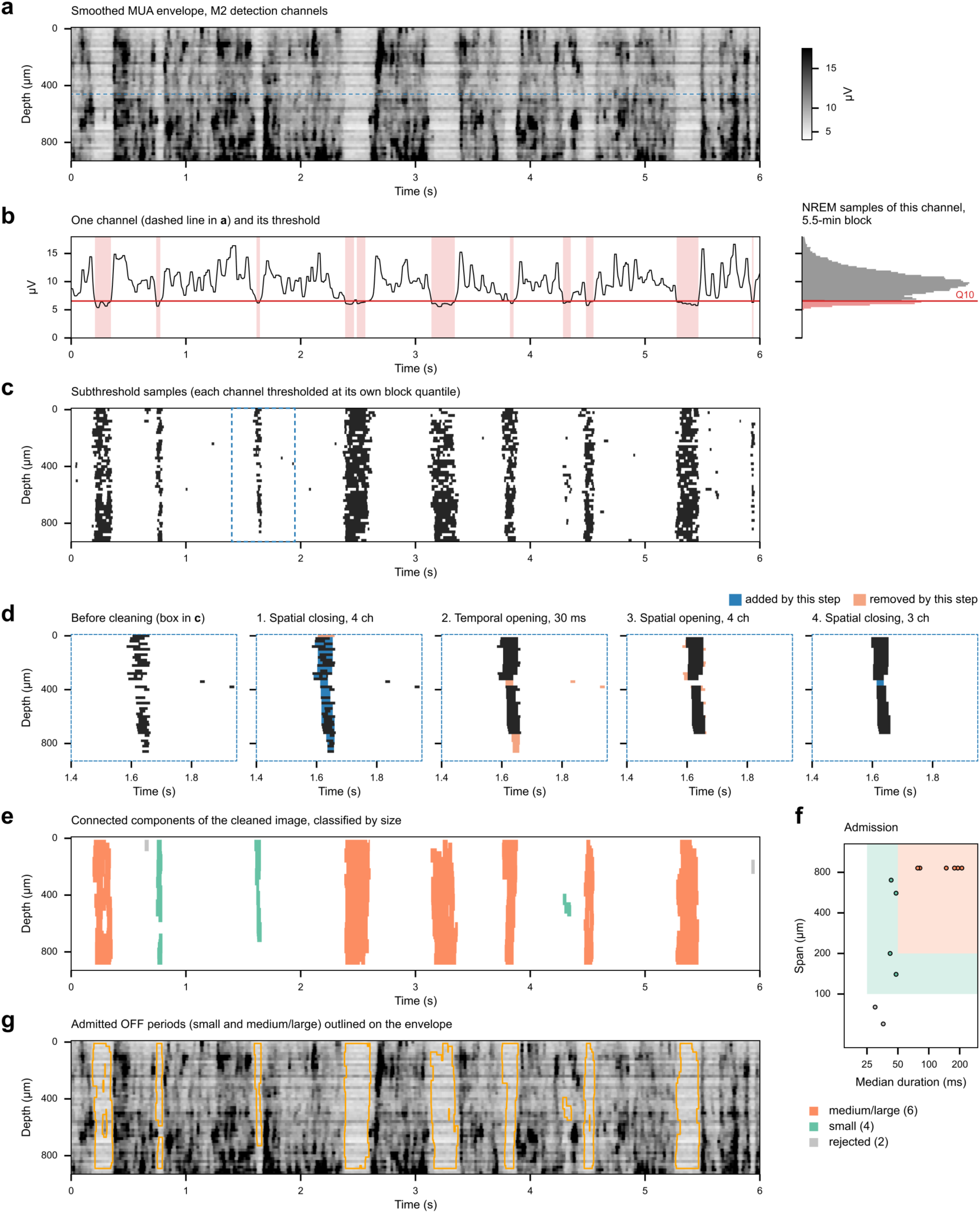
Morphological OFF period detection. A 6-s segment of early recovery NREM sleep from secondary motor cortex (M2) of one subject. Depth is measured from the most superficial detection channel (47 channels, 20-µm pitch). **a,** smoothed MUA envelope (see Methods). Darker shades indicate a higher envelope, as in Fig. 1c. The dashed line marks the channel shown in b. **b,** left, the envelope of that channel and its detection threshold (red); shaded intervals are subthreshold. Right, the distribution of all NREM samples of this channel in the ∼5.5-min processing block that contains the segment. The threshold is the q-th quantile of this distribution (here q = 0.10, the NREM quantile for this structure), so 10% of the samples (red) lie below it. The threshold is recomputed in the same way for every channel and every block. **c,** binary image of subthreshold samples, with every channel thresholded at its own block quantile. The dashed box marks the window enlarged in d. **d,** the four morphological operations applied to the binary image, in order: spatial closing across 4 channels (80 µm), temporal opening of 15 samples (30 ms), spatial opening across 4 channels (80 µm), and spatial closing across 3 channels (60 µm). In panels 1–4, blue pixels were added and orange pixels removed by that operation. **e,** connected components of the cleaned image, colored by whether they pass the relaxed criterion only (teal; small OFF periods), the stringent criterion (orange; medium/large OFF periods), or are rejected (grey), as in f . **f,** each component in e plotted by its median duration and span (log axes). The total-duration requirement of each criterion (at least 30 or 60 ms) is not drawn; in this example it changes no classification. **g,** the admitted OFF periods (small and medium/large) outlined in orange on the envelope from a, as in Fig. 1c.

**Supplementary Figure 2.**
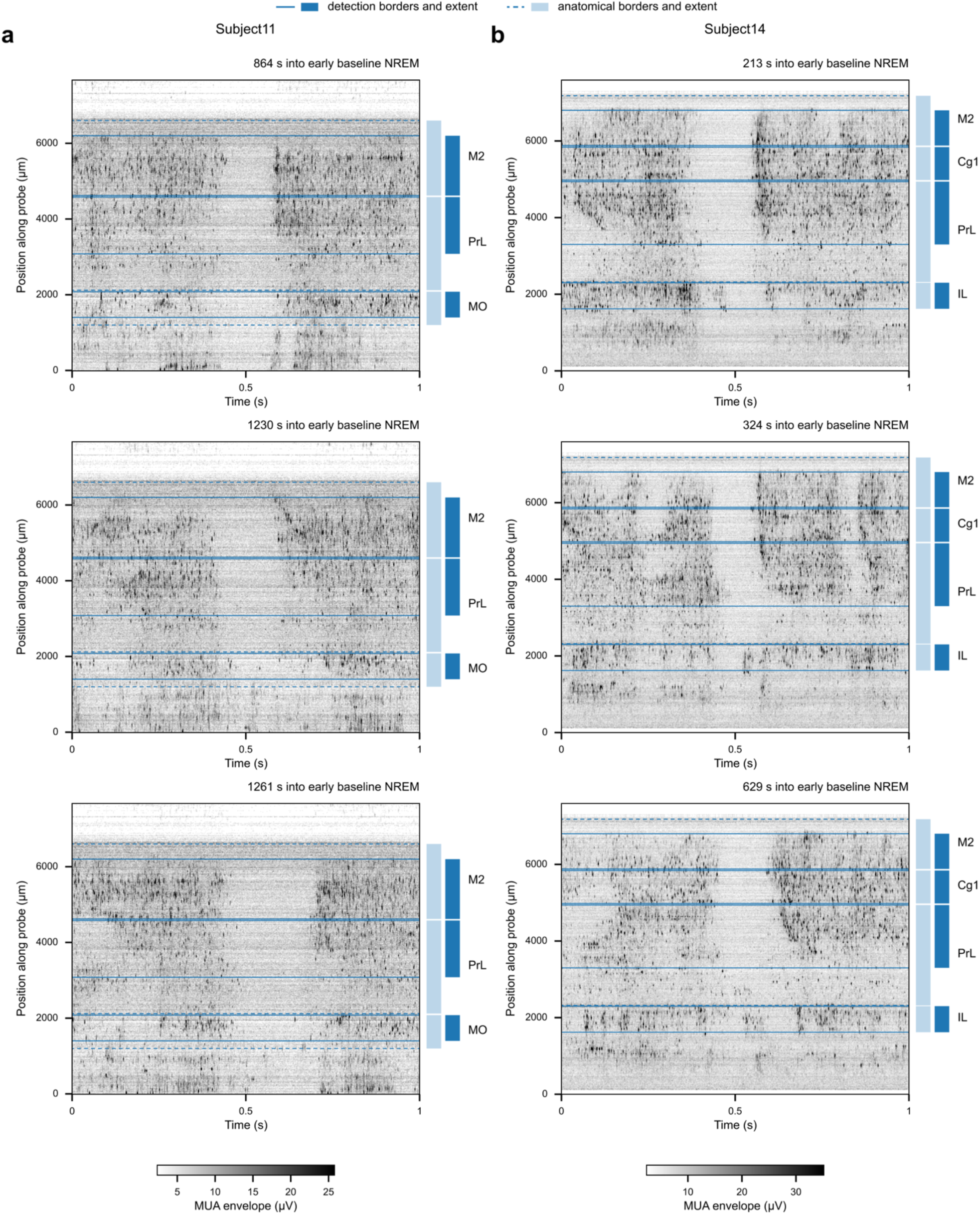
Detection borders and anatomical borders on two example probes. Each panel shows 1 s of the MUA envelope (as in Fig. 1c) on every channel of one probe during early baseline NREM sleep. **a,** Subject11 (M2, PrL and MO). Detection was not performed on the structure below MO, even though bistability is evident, because it could not be confidently assigned to a more specific structure than “ventral telencephalon” (e.g. dorsal tenia tecta vs basal forebrain). **b,** Subject14 (M2, Cg1, PrL and IL). As in a, detection was not performed on the structure below IL because it could not be confidently assigned to a specific cortical structure. Vertical position is measured along the probe from its deepest recording site, with superficial channels at the top. Darker shades indicate a higher envelope. Solid lines and dark blue bars mark the detection borders of each structure (the channels used for OFF period detection); dashed lines and light blue bars mark its anatomical borders (where the two coincide, only the solid line is visible).

**Supplementary Figure 3.**
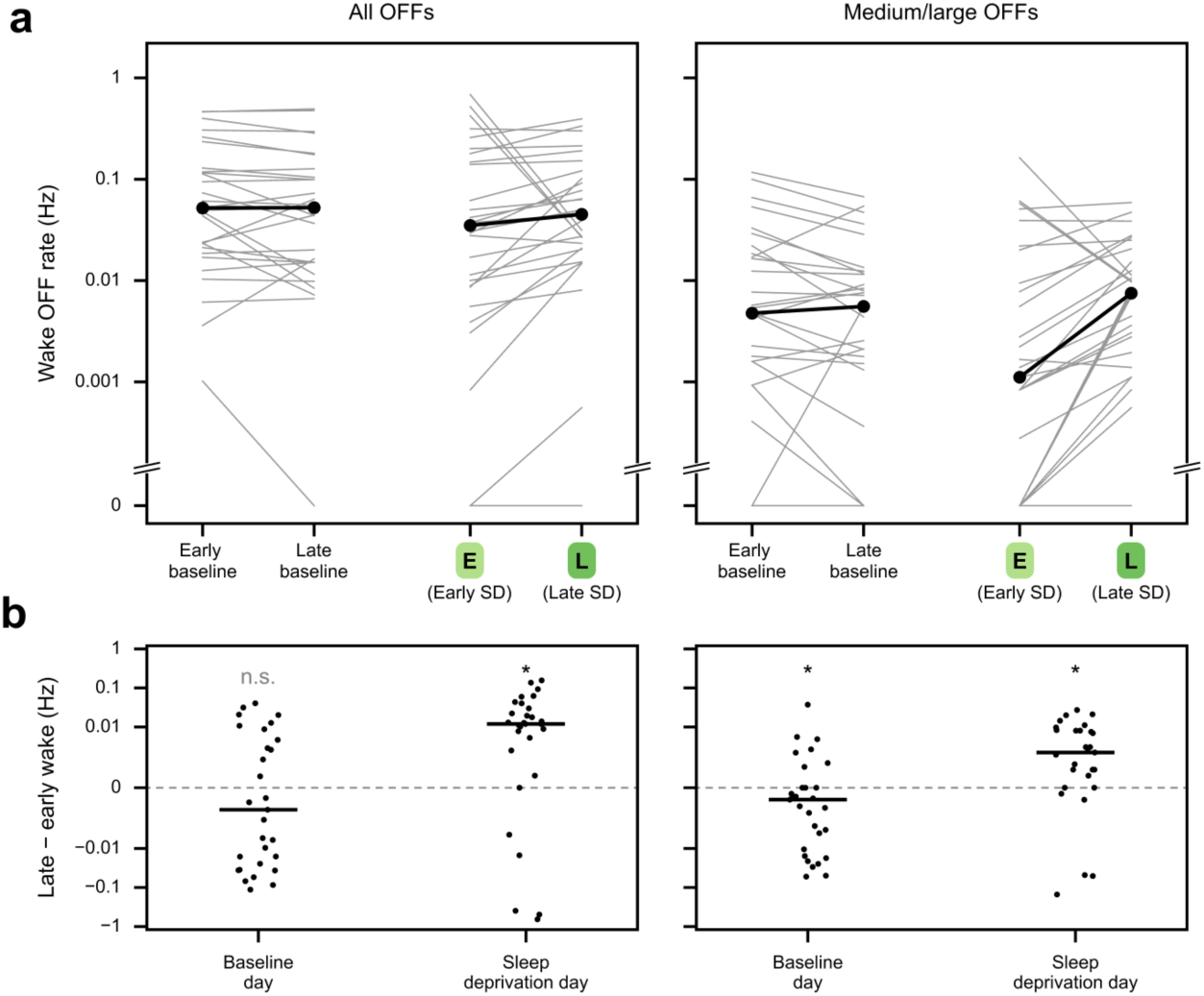
Wake OFF period rate during sleep deprivation and at the same clock times on the baseline day. For each subject-structure pair (n = 29 pairs from 15 rats), the rate of OFF periods during wake (OFF count divided by wake time) in the early and late sleep-deprivation windows (sleep deprivation day), and in wake falling within the same clock-time windows 24 h earlier (baseline day), for all OFF periods (left) and for medium/large OFF periods (right). Wake time in the baseline-day windows was a median of 18 min (early) and 31 min (late), compared with 60 min in each sleep-deprivation window. **a,** rates in each window: grey lines, individual pairs; black, medians across pairs. Pairs with no OFF periods in a window are plotted on the separate 0 row below the axis break. **b,** the late minus early difference in rate for each pair (points; horizontal lines, medians; symmetric log axis, linear between −0.001 and 0.001 Hz). Stars: two-sided Wilcoxon signed-rank tests of these differences against 0, Holm-corrected for the two days in each column (*: p < .05; n.s.: not significant).

**Supplementary Figure 4.**
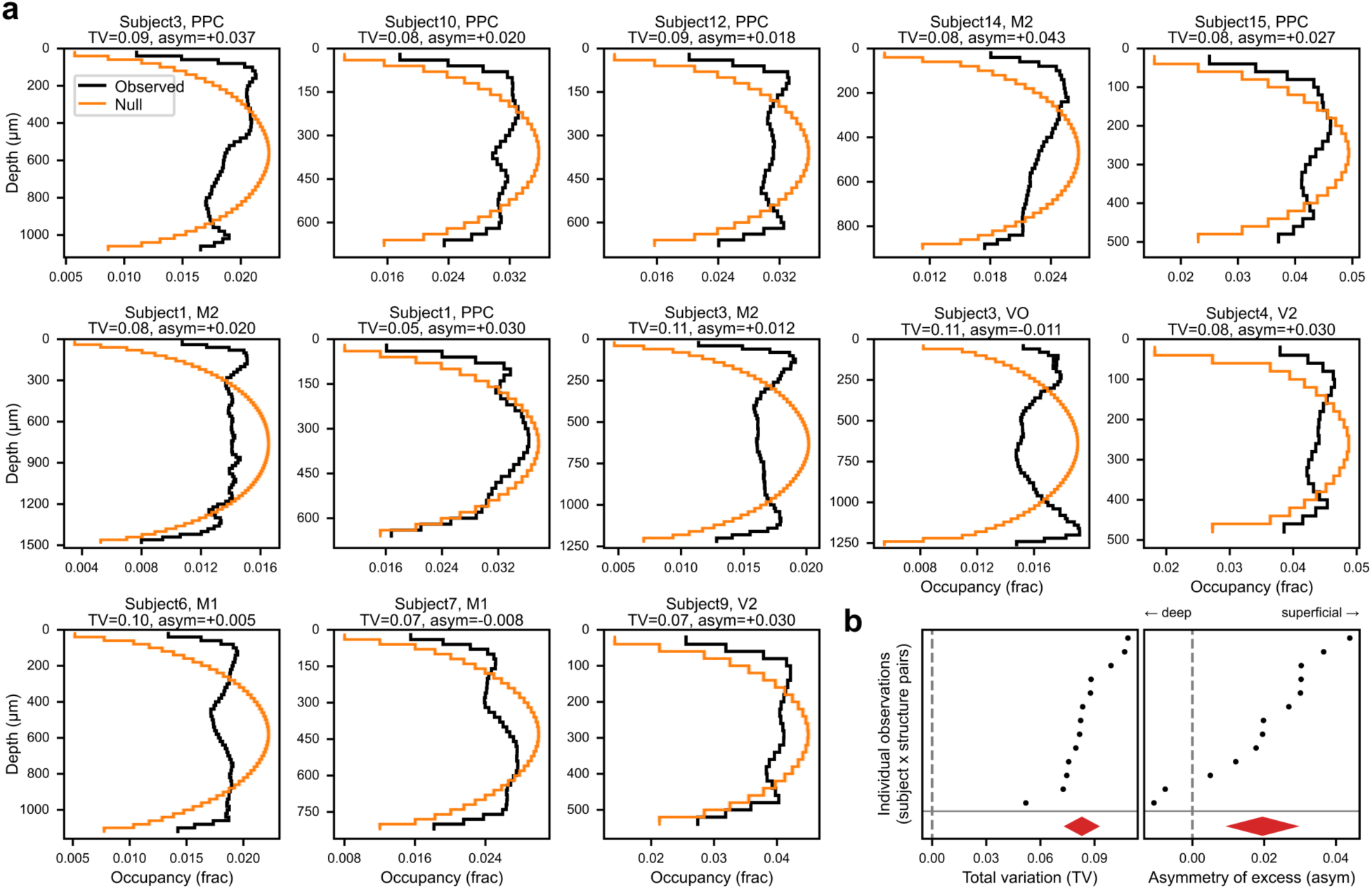
OFF-involvement by depth departs modestly but consistently from a geometry-preserving depth null. **a**, for each cortical region with sufficient span (> 500 µm) and cross-laminar probe trajectory (n=13 unique subject-structure pairs, from 10 unique subjects and 6 unique structures), the number of OFF events involving each channel as a function of channel depth (black), compared to a size- and shape-preserving depth null (orange; mean ± 2 std., band is less than the line width). The null re-drops each OFF at a uniformly random, in-bounds depth, destroying only its location along the depth axis, and assumes uniform detection precision and sensitivity across depths (see Supplementary Methods). Depth is measured from the superficial detection border of each structure, so superficial channels are at the top of each plot. TV: total variation, the fraction of OFF mass that must be relocated for the null to match the observed profile; asym: the observed-minus-null fraction of OFF mass falling in the superficial half of the span. Positive values mean the excess sits towards the cortical surface, and negative values away. **b,** group-level summary of the per-region departures in (a). Left, the TV effect for each region (one row each; rows sorted independently); right, the signed superficial–deep asymmetry of the empirical excess. Group means (diamonds) are from intercept-only linear mixed models (metric ∼ 1 + (1 | subject)), and diamond widths represent 95% confidence intervals (see Supplementary Methods; Supplementary Table S5). The reference line at 0 indicates no net superficial/deep bias.

**Supplementary Figure 5.**
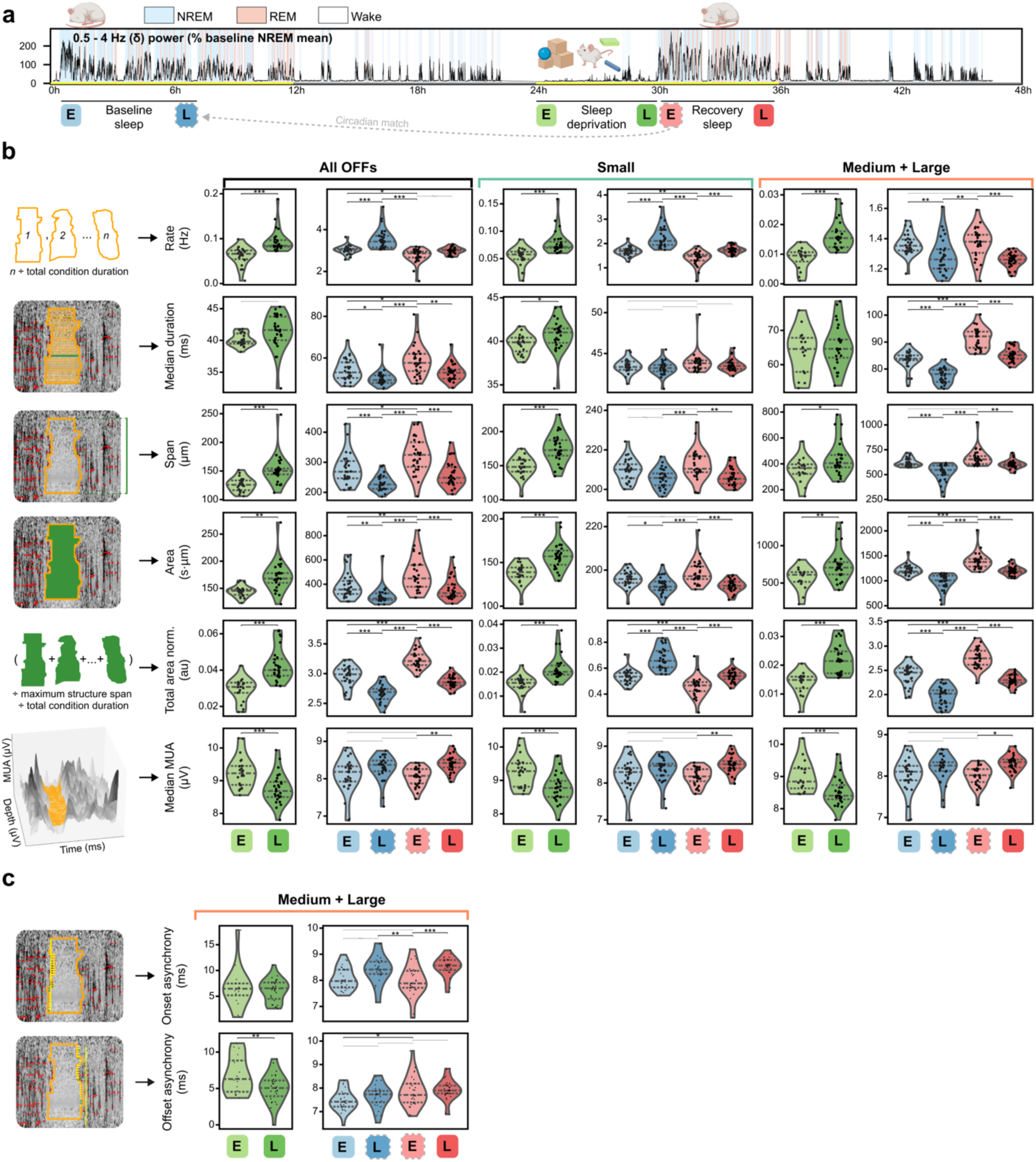
Sleep/wake changes in OFF periods by size class, and sleep/wake changes in within-structure ON-OFF transition synchrony. a, experimental protocol, as in Fig. 2a. **b,** change in temporal and spatial features of all OFF periods across experimental conditions, faceted by size class. For small OFF periods, duration and span are summarized by the mean rather than the median across OFF periods (see Supplementary Methods). “Total area norm.” is the total OFF area across the entire condition, normalized by each structure’s maximum possible span and the total condition duration, providing a measure of “total OFFness” that can be compared across subjects and structures. **c,** change in OFF onset/offset asynchrony across experimental conditions, corrected for a mechanical correlation with OFF span (see Supplementary Methods). In b-c, n=29 unique subject-structure pairs from 12 structures and 15 rats. Models, tests and significance bars are as in Fig. 2f-h. For some measures, NREM and wake conditions were fit as separate models (see Methods). Precise p-values, confidence intervals, estimates, effect sizes, and model details are provided in Supplementary Tables S0-S1.

**Supplementary Figure 6.**
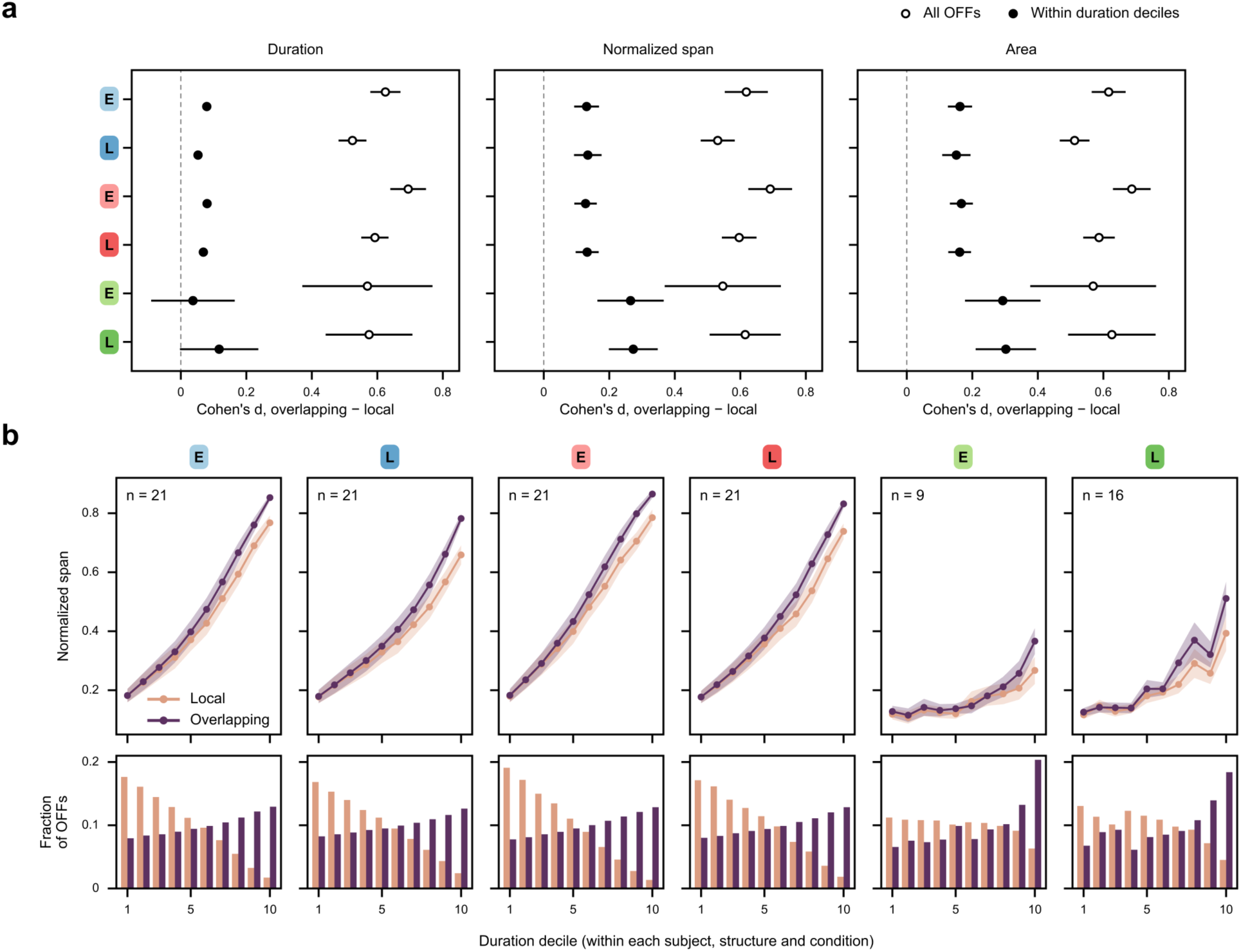
Size of overlapping and local OFF periods at matched duration (cf. Fig. 4g). Analyses use subject-structure-condition cells with at least 20 local and 20 overlapping OFF periods (n = 21 subject-structure pairs from 7 subjects and 12 structures in each NREM sleep condition; 9 pairs from 3 subjects in early and 16 pairs from 5 subjects in late sleep deprivation; conditions labeled as in Fig. 2a). Within each cell, OFF periods were split into deciles of duration. **a,** standardized difference (Cohen’s d) between overlapping and local OFF periods in duration, normalized span and area, computed from all OFF periods in each cell (open circles) or within duration deciles (filled circles: the within-decile differences averaged across deciles, weighting each decile by the product of its numbers of overlapping and local OFF periods divided by their sum, and omitting deciles with fewer than 5 OFF periods in either group; one late sleep deprivation cell had no such decile, leaving 15). Estimates are pooled across cells with a DerSimonian-Laird random-effects model; error bars, 95% confidence intervals. **b,** top, mean normalized span of local and overlapping OFF periods in each duration decile (mean ± s.e.m. across cells; deciles with fewer than 5 OFF periods in either group omitted). Bottom, the fraction of each group’s OFF periods that falls in each duration decile, averaged across cells. Median durations of the first and last deciles were 34 ms and 186–230 ms in NREM sleep conditions, and 32 ms and 88–107 ms in wake conditions.

**Supplementary Figure 7.**
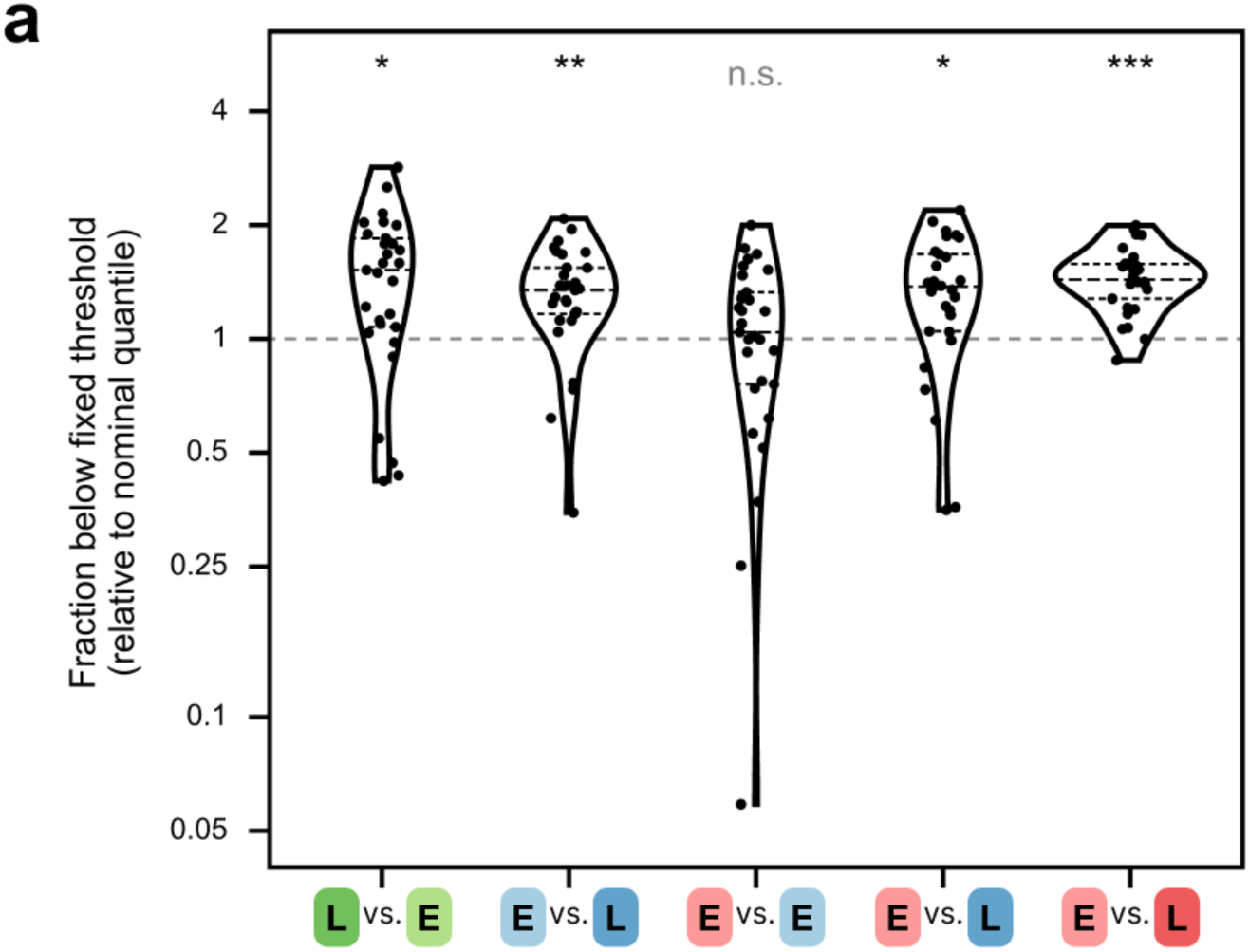
Fraction of MUA envelope samples below a fixed threshold. For each subject-structure pair (n = 29 pairs from 15 rats) and each channel, a threshold was set at the nominal quantile q of that channel’s envelope in a reference condition (the wake quantile for wake, the NREM quantile for NREM sleep), and the fraction of samples below that threshold was measured in the target condition. Points show, for each pair, the median of this fraction across channels divided by q, so a value of 1 (dashed line) means that the target condition has the same subthreshold occupancy as the reference, and values above 1 mean that more samples fall below the fixed threshold in the target condition. Comparisons are labeled target vs. reference, with conditions labeled as in Fig. 2a: from left to right, late vs. early sleep deprivation (wake); early baseline vs. circadian-matched baseline, early recovery vs. early baseline, early recovery vs. circadian-matched baseline, and early recovery vs. late recovery (NREM sleep). Violins show the distribution across pairs (dashed lines, median and quartiles; log scale). Stars: two-sided Wilcoxon signed-rank tests of the log of this ratio against 0, Holm-corrected across the five comparisons (*: p < .05; **: p < .01; ***: p < .001; n.s.: not significant).

